# The Nucleome Data Bank: Web-based Resources to Simulate and Analyze the Three-Dimensional Genome

**DOI:** 10.1101/2019.12.20.885145

**Authors:** Vinícius G. Contessoto, Ryan R. Cheng, Arya Hajitaheri, Esteban Dodero-Rojas, Matheus F. Mello, Erez Lieberman-Aiden, Peter G. Wolynes, Michele Di Pierro, José N. Onuchic

## Abstract

We introduce the Nucleome Data Bank, a web-based platform to simulate and analyze the three-dimensional organization of genomes. The Nucleome Data Bank enables physics-based simulation of chromosomal structural dynamics through the MEGABASE + MiChroM computational pipeline. The input of the pipeline consists of epigenetic information sourced from the Encode database; the output consists of the trajectories of chromosomal motions that accurately predict Hi-C and FISH data, as well as multiple observations of chromosomal dynamics *in vivo*. As an intermediate step, users can also generate chromosomal sub-compartment annotations directly from the same epigenetic input, without the use of any DNA-DNA proximity ligation data. Additionally, the Nucleome Data Bank freely hosts both experimental and computational structural genomics data. Besides being able to perform their own genome simulations and download the hosted data, users can also analyze and visualize the same data through custom-designed web-based tools. In particular, the one-dimensional genetic and epigenetic data can be overlaid onto accurate three-dimensional structures of chromosomes, to study the spatial distribution of genetic and epigenetic features. The Nucleome Data Bank aims to be a shared resource to biologists, biophysicists, and all genome scientists. The Nucleome Data Bank (NDB) is available at https://ndb.rice.edu.

## Introduction

The genome, particularly the human genome, is the object of investigation of an incredibly large and diverse community of scientists. Since the inception of the Human Genome Project,^1^ freely accessible databases have been a constant characteristic of the field of genomics, allowing the broad availability of curated experimental data that have contribute powerfully to the significant scientific and medical developments in the last 20 years. The first databases in genomic science cataloged DNA sequences, along with annotations of the functional loci in those sequences. Subsequently, it becomes clear that there is a need to collect not only information about DNA sequences but also information regarding the histone modifications and transcription factor binding sites that help to regulate genes.

We term all such additions to the bare DNA sequence as being epigenetic marks. The bare DNA sequences with the epigenetic marks provide one-dimensional descriptions of the genome that we currently believe contains much of the information needed to describe genome organization.

In the last few years, a new layer of complexity in the way the genome stores information has been appreciated; the organization of the genome in the three spatial dimensions.^2–5^ Approximately 2 meters of DNA decorated with proteins and RNA, collectively called chromatin, is contained within the nucleus of human cells. The eukaryotic nucleus is only a few microns across, requiring the genome to take on a complex structure to store so much information. On the scale of tens of nanometers, a great deal is known about chromatin structure. On this scale, we know most of the DNA is wrapped around histone protein complexes forming a string of nucleosomes. On larger length scales, the structural and dynamical organization of genes, chromosomes, and nuclei are less well understood and the subject of intense investigation. A key difficulty is that on these larger scales, in three dimensions, chromosome structures fluctuate and must be described as structural ensembles. These ensembles do have persistent structural characteristics that require storing and cataloging in databases. Creating such a catalog requires novel extensions of the way the one- and three-dimensional data are described and stored. At larger length scales, chromatin structure is specific to tissues and varies between cellular phenotypes. Clearly, the structural organization of chromosomes influences gene regulation and differentiation. Indeed, some variations in the three-dimensional structural organization of genes have been associated with diseases during embryonic development and cancer.^6–9^ The analysis of genomic architectures opens a new chapter in structural biology. The relationship between genome structure and genomic activity reminds us of the relationship between structure and function in proteins, which itself is a complex issue requiring an ensemble or landscape description.^10,11^ The relationship between the one-dimensional sequences of DNA with their attached epigenetic factors and the three-dimensional organization of chromosomes resembles the problem of protein folding, where the amino acid sequences dictate the three-dimensional fold of globular and membrane proteins^12^ and also influences the structural ensemble of the so-called intrinsically disordered proteins. Detailed knowledge of the three-dimensional organization of the genome and how it maps on to cellular phenotypes should help us understand how the genome is maintained and regulated, and shed light on important aspects of disease.

Here, we introduce a new web-based platform that aims to be a resource for biologists, biophysicists, and other genome scientists studying and using the three-dimensional genome. We have named our platform the Nucleome Data Bank in analogy with the Protein Data Bank,^13^ a data repository whose contribution to the progress of structural protein biology cannot be questioned.

### The Nucleome Data Bank

The Nucleome Data Bank is a repository of both structural genomic data and powerful computational tools that enable physics-based simulation of chromosomal structural dynamics.

The Nucleome Data Bank is offered as a shared web-platform that freely hosts data about the spatial organization of the genome by facilitating the analysis and visualization of the three-dimensional chromosomes through custom web-based tools.

Unlike folded proteins which have relatively small but non-negligible fluctuation, chromosomes have highly heterogeneous architecture and must be represented by ensembles of structures as opposed to one single, average structure.^14–16^ As a consequence, ensembles of 3D structures are the primary focus of the data bank (Figure 1).

**Figure 1:**
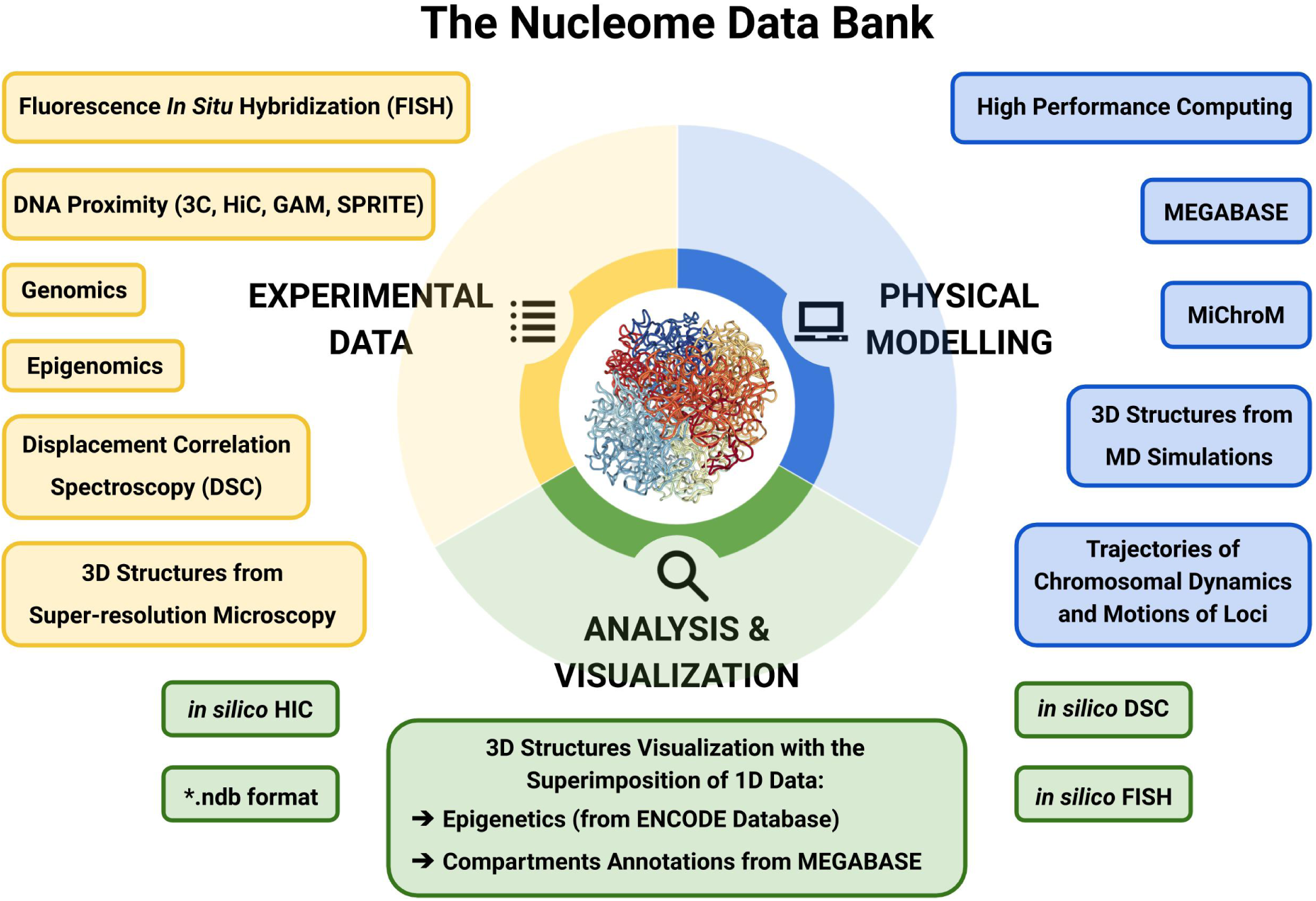
The Nucleome Data Bank (NDB) focuses on genomic 3D structural ensembles. All experiments probing genome architecture can ultimately be reconnected to these ensembles. Because of this, 3D ensembles provide a unified framework for the interpretation of all types of experimental observation and constitute the central link connecting experiments that would otherwise be difficult to compare. The NDB is a repository of web-based computational tools enabling predictive physical modeling of the three-dimensional architecture of genomes, analysis and visualization of the chromosomal structures. The 3D structures of chromosomes and their motions are predicted using the MEGABASE+MiChroM pipeline through Molecular Dynamics simulations; these simulations are typically carried on using High Performance Computing (HPC) resources. Experimental data are used both as input and validation data sets to the MEGABASE+MiChroM pipeline, which is outlined in figure 2. The physical simulations achieve specificity using as input information about the epigenetic marking patterns of specific chromosomes in specific cells. The chromosomal structural ensembles resulting from simulations can then be used to generate *in silico* Hi-C maps, as well as *in silico* Displacement Correlation Spectroscopy and FISH. Additionally, the NDB freely host genomic structural ensembles of both computational origin, as the ones from the MEGABASE+MiChroM or similar pipeline, or experimental origin, as the structures obtained from super-resolution microscopy. The 3D structure data are available in the newly defined .ndb file format. The 3D visualization of chromosome structures from simulations is also available in the NDB server. In the NDB visualization scheme, 1D experimental signal from NIH-ENCODE can be overlaid onto chromosome 3D structures to study their spatial features.

**Figure 2:**
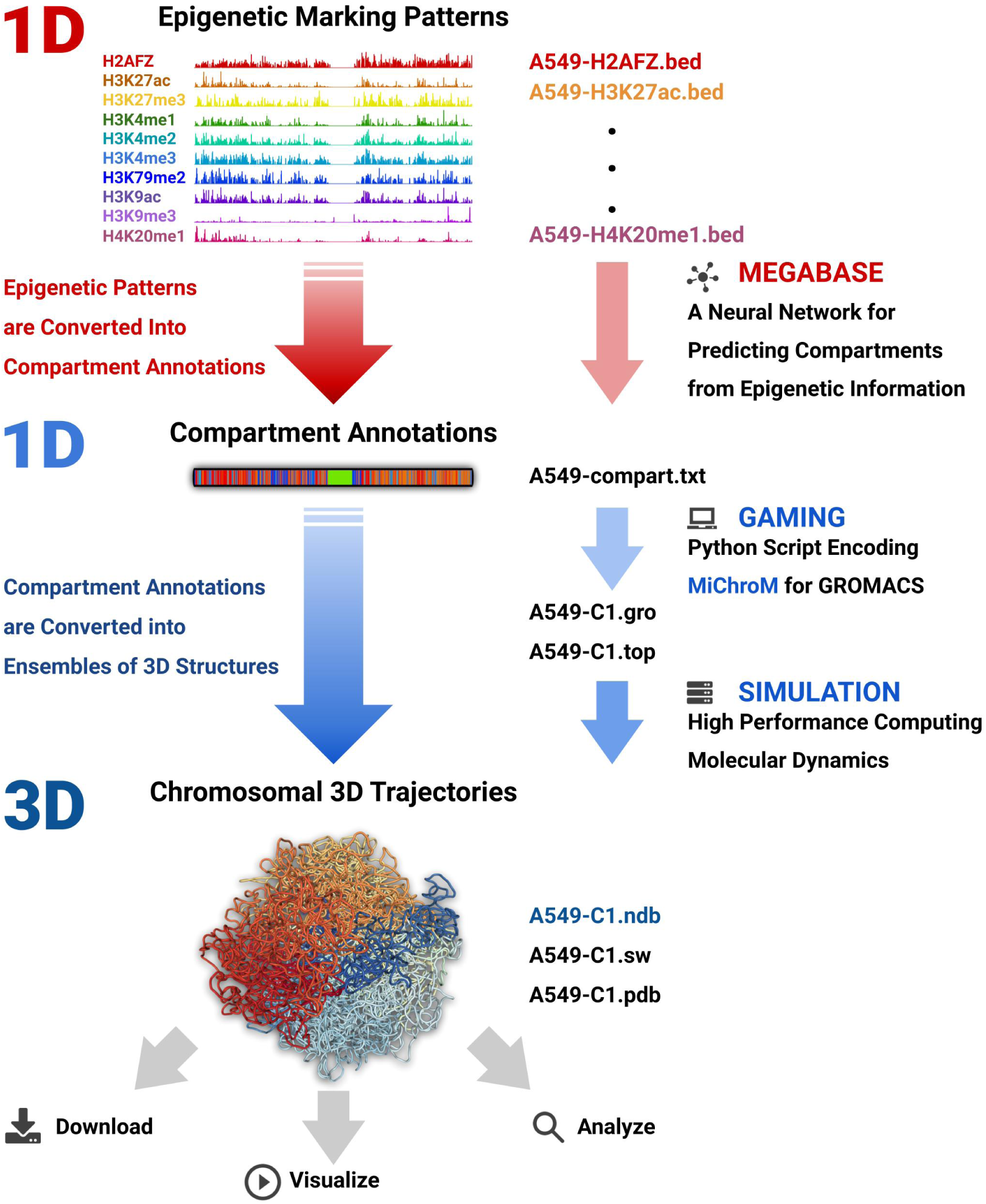
The MEGABASE+MiChroM Pipeline and its implementation. First, Chromatin Immuno-precipitation sequencing (ChIP-seq) assays about the epigenetic markings of a cell type are obtained from the NIH ENCODE Project database. From these data, the neural network MEGABASE generates annotations for the chromosomal compartments and sub-compartments. These annotations are specific to the cell line initially selected and are in turn used to model the chromosome(s) under investigation according to the Minimal Chromatin Model (MiChroM); a physical model for chromatin folding based on motor activity and micro phase-separation. Molecular dynamics simulation are then performed to produce an ensemble of chromosomal 3D structures. On the right-hand side, figure also shows the file formats involved at every step of the pipeline. Finally, the 3D chromosomal trajectories generated from these physical simulation can be used to generate a variety of predictions, such us *in silico* Hi-C maps as well as *in silico* FISH. All the structural ensembles generated so far are already stored on the NDB server, where they are available for visualization and download.

3D structural ensembles are the richest and most complete data sets to describe the genome organization; all experiments probing genome architecture can ultimately be reconnected to these ensembles. Many different experimental assays have already been explained and predicted using structural data; as such, 3D ensembles constitute a central link connecting experiments of very different sorts.

Clearly, if the structural ensemble correctly reproduces the statistical distribution of the chromosomes within cells, then Hi-C^2,17^ and other DNA proximity assays^18–23^ can be inferred by measuring the frequency of genomic contacts in the ensemble.

Similarly, the distribution of distances monitored by Fluorescence In-Situ Hybridization^12,24^ can also be easily compared with the predicted distribution of distances in the chromosomal structure ensemble. Recently, highly multiplexed FISH, in combination with super-resolution microscopy has allowed direct visualization of chromosomal segments. ^14,15^ These images can now be compared with their counterparts obtained from computational modeling to characterize the motion of chromosomal loci. The dynamics of fluorescently labeled loci has been followed in time.^25,26^ The heterogeneity of the motions of different loci, their diffusive properties, timescales of motions, and the correlations between the motions of different loci have already been analyzed with the fundamental support of Molecular Dynamics simulations (MD) of entire chromosomes. ^27^

The Nucleome Data Bank hosts both experimental and computational structural data. The information is organized by organism, cell type, chromosomal number, and when available, homologous copy. The information about the positioning of the different loci is offered in a newly defined file format which is described in detail in a dedicated section below. The new format, with extension .ndb, beside other information, provide a sampled set of structures from the ensemble for a segment of DNA or an entire chromosome. Each structure is a set of xyz coordinates that represents the conformation of the DNA polymer in three dimensions at a specific instant. When the set of structures has been recorded as a time series, then the set represents a dynamical trajectory of chromosomal motions.

At the moment of writing, the Nucleome Data Bank stores structural data about 8 human cell lines, accounting for over 11 millions of individual chromosomal structures. The vast majority of these data are trajectories sampled by MD simulations. The currently limited amount of laboratory experimental data obtained through DNA tracing is also accessible through the same platform. In the next few years we plan to continuously expand the nucleome data repository, and we encourage the authors of newly published manuscripts to share their structural data through the Nucleome Data Bank.

Beside making available all of these structural data the Nucleome Data Bank web interface allows easy visualization of the structural ensembles. The interface allows users to scroll through the structures, manipulate them in 3D by rotations and translations, zooming in and out. The interface also allows users to play movies of chromosomal dynamics in addition to providing images as single snapshots.

The entire database from the NIH Encode^28,29^ project can also be overlaid on these structures. Once the cell line and the chromosome of interest have been selected, the user can choose a one-dimensional experimental track which can be superimposed on the three-dimensional structure of the very same chromosome.

For the first time, through the Nucleome Data Bank, it is now possible to analyze and visualize the cell-specific 3D spatial conformation of a massive amount of data about DNA binding, methylation, accessibility, transcription obtained from a wide range of different experimental assays (e.g. ChIP-seq, RNA-seq, Dnase-seq, ATAC-seq).

In addition to all of the functionalities listed above, the Nucleome Data Bank makes available to the broader community of scientists a series of computational tools developed at the Center for Theoretical Biological Physics (a Frontiers of Physics Center sponsored by the National Science Foundation). In particular, the MEGABASE+MiChroM pipeline, allows for the predictive physical simulations of the chromosomal structural ensemble and chromosomal dynamics, starting from the input of only epigenetic marking data.

### Predicting Genome organization in time and space: the MEGABASE+MiChroM computational pipeline

The MEGABASE+MiChroM pipeline enables the study of the structural ensembles and motion of chromosomes through Molecular Dynamics simulations. ^30^ The input of the pipeline consists of epigenetic information sourced from the Encode database; the output consists of trajectories of chromosomal dynamics. The two methods MEGABASE^12^ (short for Maximum Entropy Genomic Annotation from Biomarkers Associated to Structural Ensembles) and MiChroM^17^ (short for Minimal Chromatin Model) together with their implementation have been extensively validated and have been shown to produce highly accurate predictions about the global architecture and the motions of labeled loci in human chromosomes.^27^

The Minimal Chromatin Model is an energy landscape model for chromatin 3D structure at 50kb resolution. This model combines the energy function characteristic of a connected polymer chain with additional interaction terms governing compartment formation, as well as modeling the coiling effects of the proteome involved in chromatin organization. The effects of molecular motors acting along the chromatin polymer are modeled by a data-driven, translationally-invariant energy term dubbed the Ideal Chromosome. ^17,31^ This energy term is currently taken to be universal and is not changed across chromosomes or cell lines. The Ideal Chromosome potential does however change in shape according to the cell cycle, this feature accounts for different levels of motor activity in different cell phases.

MiChroM introduced the idea that the compartmentalization seen in Hi-C maps arises from microphase separation of chromatin segments having different biochemical properties; using this idea, the compartmentalization patterns found in Hi-C maps can be transformed into ensembles of 3D models of genome structure at 50-kb resolution. Clearly, these patterns are not universal, as every chromosome and every tissue or cell line from the same organism exhibits its own compartmentalization pattern.^32^ Similarly, the distribution of CCCTC-binding factors (CTCF) along DNA is specific to distinct chromosomes, and therefore, so are the resulting looping patterns, also observed in Hi-C maps.^4,33^

In MiChroM the 3D interphase architecture of a chromosome is encoded in its one-dimensional sequence of compartment annotations and in the placement of its stable loops in the same way 3D protein structures are determined by the one dimensional sequence of amino acids along with the position of disulfide bonds.

In contrast to the situation for proteins, however, the sequence provided by the compartment annotations is not fixed on the DNA polymer (and thus it is not directly related to the DNA sequence alone) but changes according to the tissue, cell type, and phase in the cell cycle. We have already shown that the compartment annotations can be accurately predicted based exclusively on epigenetic marking patterns. These patterns seem to orchestrate ultimately the global architecture of chromosomes and possibly collaborate in modulating gene expression in different cell types. The free parameters in the MiChroM energy function were optimized so as to reproduce the Hi-C map of chromosome 10 of GM12878 cells at 50kb resolution. The MiChroM energy function contains 27 parameters (see SI^17^), which have not been recalibrated from the original publication.^17^ Now that this energy function has been learned, it is possible to perform molecular dynamics simulations of chromatin using as input the classification of loci into chromatin types and the location of loops.

The structural ensemble for a chromosome that results from following the molecular dynamics with the MiChroM energy function closely reproduces many independent experimental data sets. Using the structural ensembles obtained *de novo* through MD simulations, the pipeline predicts with great accuracy the Hi-C maps of multiple human cell lines,^32^ as well as three-dimensional imaging data obtained from fluorescence in situ hybridization (FISH). The chromosomal trajectories obtained through MD simulations can also be used to study the motions of chromosomes. This is a distinct advantage of using physics-based simulations as opposed to simpler approaches based on statistics or information theory that do not employ a macromolecular representation. Simulations of the dynamics of interphase chromatin generated by MiChroM *in silico* display spatial coherence, viscoelasticity, and subdiffusivity of the motions chromosomal loci.^27^ The very same behavior was observed in numerous FISH experiments that track temporally in three dimensions the motions of labeled chromosomal loci *in vivo*.^25,26,34^ MiChroM introduces a sequence to structure relationship between compartment annotations and chromosomal architecture. For truly predictive physical simulations of the genome the last problem standing is to generate the compartment annotations without the use of Hi-C or any other structural experimental assay; to accomplish this goal we developed MEGABASE.^12^

Besides enabling *de novo* simulations, MEGABASE solves a very practical problem. While epigenetic marking data are relatively easy to get, generating sub-compartment annotations from Hi-C requires extremely deep sequencing making it very expensive. As a result, at the moment of writing, experimental sub-compartment annotations have been generated only for one cell line, the human lymphoblastoids GM12878.^4^ Generating sub-compartment annotations using MEGABASE is, in contrast, cheap and easy, making it a very useful computational tool, independently from its ability to carry out physical simulations of the chromosomal structural ensemble. Given the scarce availability of direct experimental compartment annotations, we have generated compartment and sub-compartment annotations for all the cell lines found in the ENCODE database, with the exception of a few cell lines that still have insufficient marking data; these annotations, for 216 cell lines, can be found in the NDB data repository. Additionally, we allow users to upload their own ChIP-seq data to the NDB server in order to generate the same annotations for their own non-standard cell lines.

MEGABASE is a neural network that uses epigenetic information in the form of ChIP-seq signals to predict genomic sub-compartment annotations without the need to carry out any DNA-DNA proximity ligation assays. The associations between biochemical markers and structural compartmentalization features have been learned using the experimental compartment annotations for the cell line GM12878 mentioned above. As we showed, the connection between structure and epigenetics is a transferable feature of genome architecture, and the neural network in MEGABASE is, therefore, able to predict compartments in a variety of other human cell lines.^32^

By combining MEGABASE and MiChroM it is possible to perform predictive simulation of chromosomes from input epigenetic data. First MEGABASE is used to convert multiple 1D epigenetic signals into a single 1D sequence of compartment annotations; the NDB server performs this operation for the user. Then using the MiChroM energy function it becomes possible to perform realistic physical simulations of the chromosomal dynamics.

It is important to stress that, while other methods^35–40^ seek to predict DNA-DNA ligation data from ChIP-seq, we do not specifically pursue this aim; instead, we seek to predict directly the 3D structural ensembles of chromosomes. *In silico* ligation data are generated from the 3D structural ensembles only for validation purpose. Because these 2D ligation data are easily generated by coarsening the 3D structural data, we do not store these data sets on the NDB. Consistently, MEGABASE was not developed to *exactly* reproduce the compartment annotation from DNA-DNA ligation, as this would constitute detrimental over fitting. Segments of chromatin marked as to belong to one compartment can indeed find themselves spatially segregated in a different compartment, just because they happen to be tethered to large amounts of differently marked chromatin. Divergences between the MEGABASE biochemical compartment annotations and the structural annotations from ligation data are expected; these divergences are often resolved by generating the structural compartment annotations from the *in silico* ligation data resulting from 3D simulations.

Performing realistic physical simulations of chromosomal dynamics is computationally intensive and requires the use of dedicated High Performance Computing (HPC) resources; for this reason it is not possible for the NDB server itself to perform simulations for the user. Instead, the NDB server creates a suite of input files for the Molecular Dynamics package GROMACS,^41^ which is a widely used, free, open source software package able to perform efficient simulations in large HPC clusters. The files generated by the NDB server contain all the information needed to produce chromosome-specific trajectories according to the input ChIP-seq experimental signals and according to the MiChroM model. GROMACS is used without needing of any modification. In the next paragraphs we will discuss details of the implementation and a test case in which we illustrate all the steps to be followed in order to generate the structural ensembles of a human cell line directly.

### Design and Implementation

The Nucleome Data Bank (NDB) is hosted by Rice University. It is available at the URL https://ndb.rice.edu. The NDB uses Django Python Web Framework version 2.1.4 and HyperText Transfer Protocol Secure protocol (HTTPS). Each job submitted to NDB generates a Universally Unique IDentifier (UUID) to ensure the user’s privacy. The back-end tools involve a combination of UNIX Shell and Python scripts. The front-end is a web interface designed to warrant an easy and intuitive user experience. NDB organizes stored 3D chromosomal structures by cell line, year, journal, and first author name. Structures from many cell lines account for many Terabytes of data, with single chromosomal trajectory files being typically of the order of Gigabytes in size. Users can download these files compressed as .zip archives. 3D structural ensembles are stored in the .ndb format but can easily be converted to spacewalk or pdb format with command-line scripts that are provided.

The first step in the simulation pipeline is MEGABASE; The tab marked as “software” redirects the users to this tool. MEGABASE converts ChIP-seq signals into compartment and sub-compartment annotations. The ChIP-seq tracks of histone modifications used in MEGABASE are H2AFZ, H3K27ac, H3K27me3, H3K4me1, H3K4me2, H3K4me3, H3K79me2, H3K9ac, H3K9me3, and H4K20me1. MEGABASE can use ChIP-seq information sourced from ENCODE database or corresponding information provided by the user. All the available information on ENCODE has been pre-processed, with compartment annotations for over 200 cell lines already available for the user to download. Experimental ChIP-seq signals are stored in the standard .bed format. .bed files are processed by integrating the signal over a 50kb bin to match the resolution of the physical simulations. ^17,30^ The ChIP-seq tracks are processed using Shell scripts while the signal integration was coded in Python. The integrated ChIP-seq signals are then used as input to the MEGABASE neural network to calculate the compartment annotations. This process takes approximately 15 minutes; if an email address was provided in the job submission, the user is notified upon completion. A .zip archive containing all the results is generated. The output data consist of text files containing the chromatin sub-compartment annotations (A1, A2, B1, B2, B3, and B4) for each 50kb locus of all the 22 autosomes. The MEGABASE neural network is implemented in MATLAB.^42^

Using the information about the compartment annotations obtained by MEGABASE, the NDB server then provides the user with all the necessary files to perform a molecular dynamics simulation of entire chromosomes or of segments of a chromosome. All cell lines from ENCODE have been pre-processed and their respective files are already available for the user to download. Similarly, users can also generate MD input files for their cell line by providing ChIP-seq data, compartment annotations, and loop lists. The MiChroM potential MD files are prepared for the GROMACS molecular dynamics package. The MiChroM effective energy function is implemented in GROMACS through tables. In this way, users can download and install a standard version of GROMACS package and perform chromosomal molecular dynamics simulations using only the files provided by the NDB server. The energy function is encoded in five tables with .xvg extension:

- **table_b0.xvg**: Polymer chain connectivity. Includes Finite Extensible Nonlinear Elastic (FENE) bonds together with nearest neighbors steric interactions.
- **table_b1.xvg**: Nuclear confinement mimicked by a repulsive spherical surface.
- **table_b2.xvg**: Ideal Chromosome potential.
- **table.xvg**: Chromatin type-to-type interactions together with the non nearest neighbors steric interactions.
- **tablep.xvg**: Stable chromatin loops defined by their anchor points.

The NDB server provides two other files with extensions .gro and .top:

- **\*.gro**: A coordinate file containing the initial position of the polymer in 3D space, the *xyz* coordinates for each chromosome locus at 50kb resolution. The file also contains the compartment annotations for each segment.
- **\*.top**:A file containing information about the topology of the system, i.e., polymer connectivity, and also containing information about locus-to-locus physical interactions.

A python script named GAMING (Genome Architecture MiChroM input files for Gromacs) performs all the tasks needed to produce the files listed above. The MD input files generated by GAMING are typically on the order of Megabytes in size for a single chromosome. A step-by-step tutorial page on how to perform chromosomal MD simulations is also available in the NDB documentation. These simulations are computationally very intensive and should typically be carried out by taking advantage of High Performance Computing resources. Python scripts useful for analyzing the output data of the MD simulations are also provided and explained in the tutorial page; in particular, it is possible to generate *in silico* Hi-C maps from the MD trajectories that are generated. The MD simulation output is a time series of 3D structures composed of hundreds of thousands of individual chromosomal configurations. Each configuration consists of the xyz coordinates of all simulated loci. Simulated structural ensembles for many cell lines are already available for download as are several experimental datasets. The NDB webpage also allows an intuitive visualization of structural ensembles. For the first time, the one-dimensional information in the ENCODE database can be overlaid onto a realistic 3D structure set to study its spatial features. The user can select different coloring schemes for the chromosome such as genomic index (position along the polymer), compartments (chromatin structural types), and ChIP-seq signals; other experimental data will soon be available. The visualization platform was implemented using NGL Viewer web application.^43^ The user can play a movie of a trajectory and explore the chromosome dynamics over time and save the structure on any selected frame. The coloring of the structures using Encode ChIP-seq tracks was developed in Python and JavaScript and is integrated with the NGL Viewer. The signal integration of a track follows the same strategy employed in MEGABASE. The color scale is proportional to the intensity of the integrated signal. The user can select different colors for the selected track using an RGB color pick embedded in NDB visualization page. The user can also change the color intensity by using the range bar.

### The NDB File Format

A few decades ago, the field structural biology faced the problem of defining a data format able to store and disseminate the increasing number of protein 3D structures obtained from X-ray crystallography. The same file format also needed to be sufficiently rich in information to support physical simulations of macro-molecules. The answer to both problems was the Protein Data Bank format (.pdb), which has since then become the standard file format for macro-molecular data. ^44^ Today, the field of structural genomics faces the very same challenge,^45^ with the increasing availability of experimental data about genome architecture and the development of new schemes for physical simulations of genome dynamics. Standard .pdb files do not address the specific necessities of structural genomics while no other standard file format has been established yet.

Here, we introduce the Nucleome Data Bank (.ndb) file format (Figure 3). The .ndb is a human-readable textual file format designed in continuity with the established practice of the structural biology community and with a strict analogy to its predecessor, the Protein Data Bank format. The .ndb file header, similar to .pdb, contains information about how the 3D coordinates were collected. The header could also include the title of a publication together with the authors’ names and journal. Just as for protein amino acid sequences, the header reports the sequence of compartment annotations for each segment of chromatin polymer for each chromosomal chain. The .ndb file stores one or more structures of the same molecular system in a single file. This could be used to store several experimental models as well as 3D trajectories from simulations. The body of the .ndb file contains twelve fields. The first field defines the type of nuclear element, CHROM for a segment of chromatin or DNAELE for any other biological element associated with DNA, whether that element is bound to DNA or is free to diffuse. More element types can be defined in the future and reported in this field. The second field reports the index of the nuclear element. The third field carries information about the epigenetic state of the locus; in this case the field contains the subcompartment annotations generated by MEGABASE. The entry notation follows a two-letter code representing the standard sub-compartment annotations (A1, A2, B1, B2, B3, and B4). This field also uses the two-letter code to identify DNA elements., e.g., TF for transcription factor and CH for the cohesin protein complex. The field number four contains information about genetics and describes properties of the DNA sequence, such as protein binding sites or location of promoter sequences; such as a CTCF binding site together with its orientation for example. The fifth field reports the chain identifier that labels distinct chromatin polymers or DNA elements. For example, each of the two copies of chromosome 1 could be labeled, as 1M, for the maternal copy and 1P for the paternal. Field number six is a progressive index identifying each entry within a single chain. DNA elements entries constitute their own chain composed of only one monomer. The fields seven, eight, and nine represent the 3D coordinate X, Y, and Z, expressed in the units of length defined in the header. The fields ten and eleven indicate the start and end positions of the DNA segment in units of base pairs, according to the genomic assembly reported in the header. The last field (number 12) is dedicated to experimental data sets and reports about the width of a locus’ spatial distribution. In optical experiments, the locus coordinates are determined by the middle point of a disperse distribution of fluorescence dyes; it is therefore useful to report the spread of such distribution. Finally, the .ndb file format reports about the positioning of stable chromatin loops. Stable loops are defined by the index (field number 2) of their anchors and by their nature if that is known, COHESIN or POLYCOMB for example. The.ndb file format was built to be maximally compatible with all the available software for 3D visualization, which was typically designed for .pdb files, as well as all the software designed compress .pdb files into binary formats.

**Figure 3:**
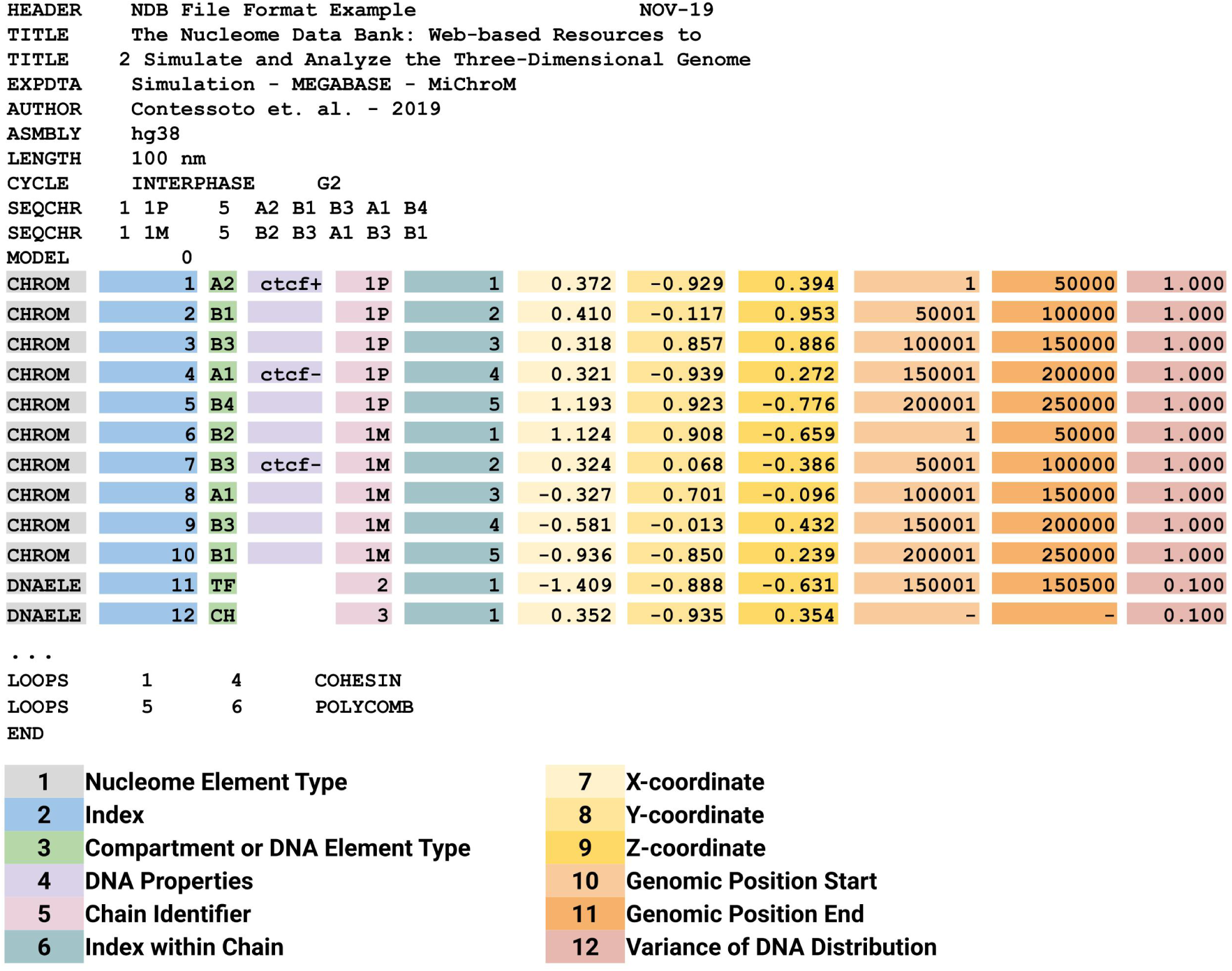
The .ndb file format. The .ndb format is designed to store data about the 3D conformation of genomes and contains enough information to be a sufficient input for physical simulations of chromosomes. The .ndb file format is inspired by the .pdb format and it is designed in strict analogy with it. The file contains twelve fields marked in figure by different colors. The first field describe the type of nuclear element: CHROM for a chromatin segment or DNAELE for any other biological element associated with DNA, whether this element is bound to DNA or free to diffuse in the nucleus. The second field is the sequential index associated to the nuclear element. The third field reports epigenetic information for the CHROM elements (chromatin type annotations in this case) or any two-letter identifier for DNAELE, TF, for a transcription factor, CH for cohesin protein complex, for example. The fourth field contains information about the DNA sequence, such as the presence of a CTCF-binding motif together with its orientation. Field number five labels the different molecular chains, e.g., 1P for the paternal copy of chromosome 1 and 1M for the maternal copy. Field number six contains a progressive index for the elements within their respective molecular chain. Fields seven, eight, and nine are the spatial coordinates X, Y, and Z of the nuclear element. Fields ten and eleven represent the start and end genomic coordinate of the locus in the case of CHROM element. For DNA-bounds DNAELE elements fields ten and eleven indicate the binding site, while a dash indicates that the element is unbound. In case of experimental data, the last field reports the variance of the spatial distribution observed for the associates locus

Other files formats for storing three-dimensional information of chromosomes have been proposed such as the spacewalk format (.spw) (http://3dg.io/spacewalk/), based on the .bed format (Browser Extensible Data). As much as the .pdb format is the standard format of the structural biology community, the .bed format is the standard format of the genomics community. While the .bed format is not well suited for molecular dynamics simulation, it is already sufficient to store the genomic position of loci together with their spatial coordinates, as resulting from DNA tracing experiments for example. In addition to the .pdb-inspired .ndb format, the Nucleome Data Bank also supports the spacewalk format (.spw). It is worth noting that the .ndb format is in fact a superset of the .spw format, with a .spw file consisting of the fields 1, 10, 11, 7, 8, and 9 of the corresponding .ndb file. Besides storing files in both formats, through the NDB web page we also provide command-line tools to convert data from .ndb format to .pdb, .spw, .gro, and vice-versa.

### A Case Study

We present here a case study in which we start from publicly available histone modification ChIP-seq signals and then generate compartment annotations. We then sample the 3D structural ensembles for all autosomes using molecular dynamics. Finally we validate the simulated ensembles by building *in silico* Hi-C maps and comparing them to the maps determined through wet lab experiments. A similar validation has already been carried out for multiple cell lines.^32^ For this case study, we will use the adenocarcinomic human alveolar basal epithelial cells (A549 cell line) for which both histone modifications ChIP-seq tracks and Hi-C maps are available. In the case of a cell line for which there are no publicly available data, users can provide their own ChIP-seq signals. For the A549 cells in this case study, all available data were already processed. The resulting files are available for download in MEGABASE page by selecting the cell line A549. The files set contains a folder for each chromosome named C1, C2, …, C22. Each chromosome was simulated using the HPC resources of Rice University. Thirty replicas of each chromosome were simulated to collect sufficient statistical sampling. Each replica simulation yielded at least 17,500 3D structures. Collectively, the number of collected structures exceeds 11 million, accounting for over 500Gb of data. These data are available for download on the NDB Data page. A fraction of the generated 3D structures is also available in the NDB Visualization page. Figure 4 shows a representative structure for each one of the 22 autosomes simulated using the MiChroM potential in GROMACS. Structures are visualized using the visualization tools of the NDB server and color-coded by using the “Color by Index” selection. The “Color by Index” option associates with each locus a color according to its genomic position from head to tail, red to blue respectively. Figure 4A contains a larger snapshot of the 3D structure of chromosome 12 also shown in the index representation.

**Figure 4:**
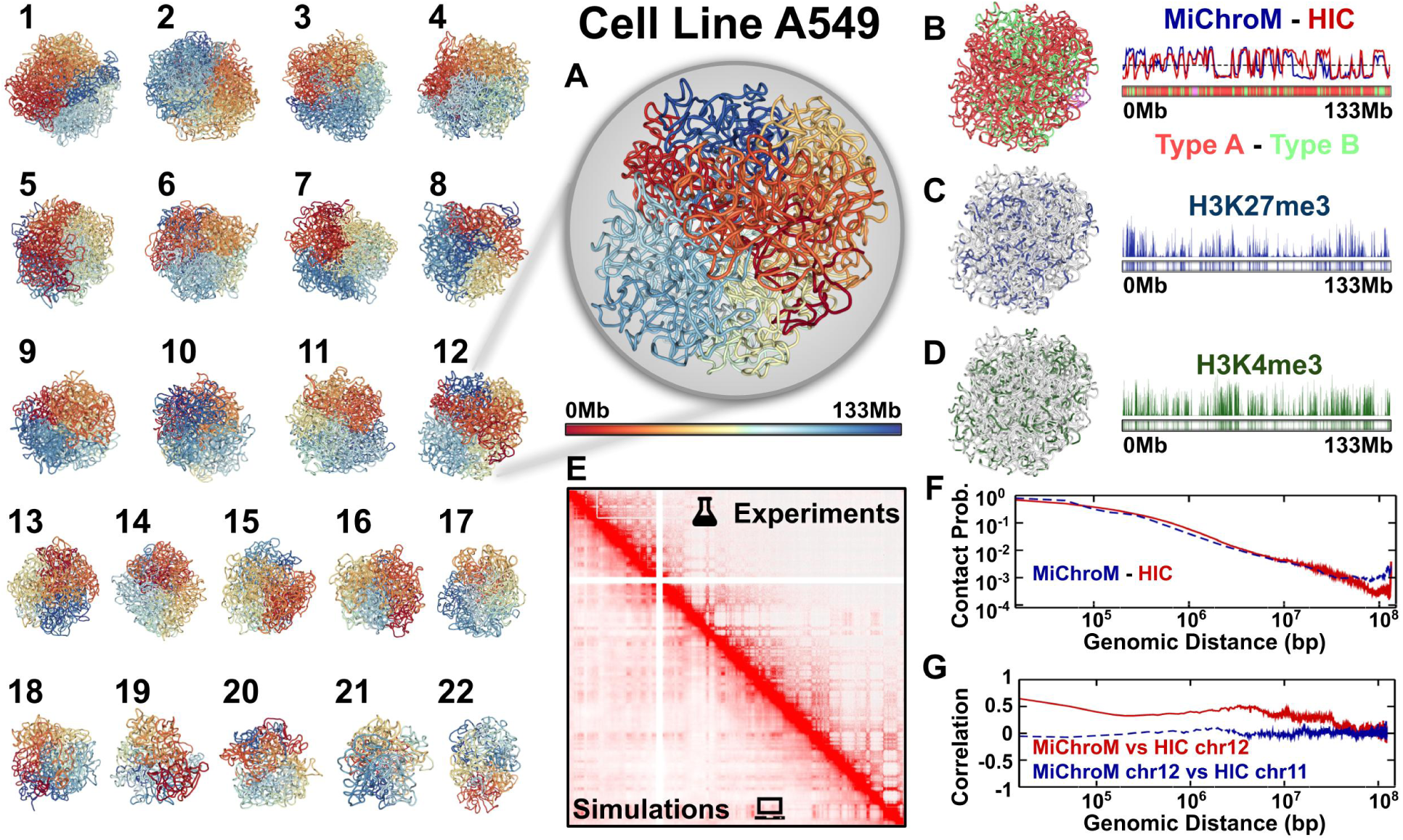
Predicting and analyzing the structural ensembles for all the autosomes of human lungs cells A549. (1-22) Representative 3D structures for all the 22 autosomes generated by Molecular Dynamics simulations according to the MiChroM potential. All the structures were obtained by using the screenshot functionality in the NDB-Visualization page. Chromosomal loci are colored according to their genomic position, from head to tail, red to blue, using the “Color-by-Index” option. (A) An enlarged view of the structure of chromosome 12. Chromosome 12 is used as an example of the analysis in all the subsequent panels. (B) The structure of chromosome 12 color coded according to the compartment annotations obtained from MEGABASE. Chromatin type A and B are shown in orange and green, respectively. Centromeric regions are instead shown in pink. compartment annotations correlate with the eigenvector of the Pearson matrix extracted from experimental and *in silico* Hi-C maps^2^. The simulated eigenvector is highly correlated with the experimentally determined one as shown on the right-hand side. (C and D) 1D ChIP-seq tracks overlaid the 3D structure of chromosome 12. Figure shows the experimental signals related to H3K27me3 and H3K4me3, in blue and green respectively. (E) Panel shows a comparison between the experimental (top) and *in silico* (bottom) Hi-C maps. The two maps are highly correlated. (F) Contact probability as a function of the genomic distance. The dashed blue line represents the curve obtained from molecular simulations using the MiChroM potential. The solid red line shows the data extracted from the experimentally determined Hi-C map. (G) Pearson’s correlation between experimental and simulated Hi-C maps of chromosome 12 as a function of the genomic distance, shown by the solid red line. As a term of comparison the dashed blue line shows the correlation between Hi-C maps of different chromosomes.

A second option is coloring the chromosome by the compartment annotations A and B, orange and green, respectively (Figure 4B). The subcompartment annotations were generated by MEGABASE, since no annotations from Hi-C are presently available for this cell line, as for many others. Coarser compartment annotations can instead be extracted from the available Hi-C maps by using the first eigenvector of the Pearson correlation matrix.^2^ A comparison between the first eigenvector extracted from the experimental and the *in silico* Hi-C maps is shown in Figure 4B. The two vectors are highly correlated (Pearson’s R = 0.72). The third option of coloring is based on the ChIP-seq signals available from the ENCODE database. Figures 4C and 4D show a structure color coded using the experimental signals of H3K27me3 and H3K4me3, in blue and green, respectively. Using the structural ensemble from physical simulations, we generate *in silico* Hi-C maps. The Python script (GAMINGplot.py) used to process the 3D structures is available for download on the tutorial page. The predicted Hi-C maps closely reproduce the experimentally determined ones. The Pearson’s R between the two maps is between 0.90 and 0.97 for all chromosomes in A549 cell line. Figure 4E shows a comparison between the experimental (top) versus *in silico* (bottom) Hi-C maps; The maps report the frequency of contact between two loci. An ensemble of 600,000 3D structures of the chromosome 12 was used to build the *in silico* Hi-C map. Chromosome simulations employing the MiChroM potential have already been extensively validated, and reproduce FISH data, as well as many dynamical observables. More coarsegrained modes of comparison also show the accuracy of the predictions. Figure 4F shows the contact probability as a function of the genomic distance. The experimental data (solid red) and simulation results (dashed blue) fall off with distance in a similar decay.

Figure 4G presents Pearson’s correlation as a function of the genomic distance between the experimental data and the computational prediction for the same chromosome. A comparison between the data obtained from experiment for two different chromosomes is also shown. We can see that the MEGABASE + MiChroM pipeline predicts features specific to the input epigenetic sequence data and not simply generic features of all chromosomes. Results for all other chromosomes are presented in the Support Information.

### Future Developments

The Nucleome Data Bank aims to be a shared resource for biologists, biophysicist and all genome scientists. The data repository as well as the computational tools available on the server are expected to be frequently updated and expanded on a rolling basis. The authors of the manuscript and developers of the Nucleome Data Bank welcome contributions from all scientists.

## Acknowledgement

This research was supported by the Center for Theoretical Biological Physics sponsored by the NSF (Grants PHY-1427654 and CHE-1614101) and by the Welch Foundation (Grant C-1792). JNO is a Cancer Prevention and Research Institute of Texas (CPRIT) Scholar in Cancer Research. PGW acknowledges the support from the D. R. Bullard-Welch Chair (Grant C-0016) at Rice University. VGC is a Robert A. Welch Postdoctoral Fellow and was also funded by FAPESP (São Paulo Research Foundation and Higher Education Personnel), and CAPES (Higher Education Personnel Improvement Coordination) Grants 2016/13998-8 and 2017/09662-7.

## Supporting Information for

**Figure S1:**
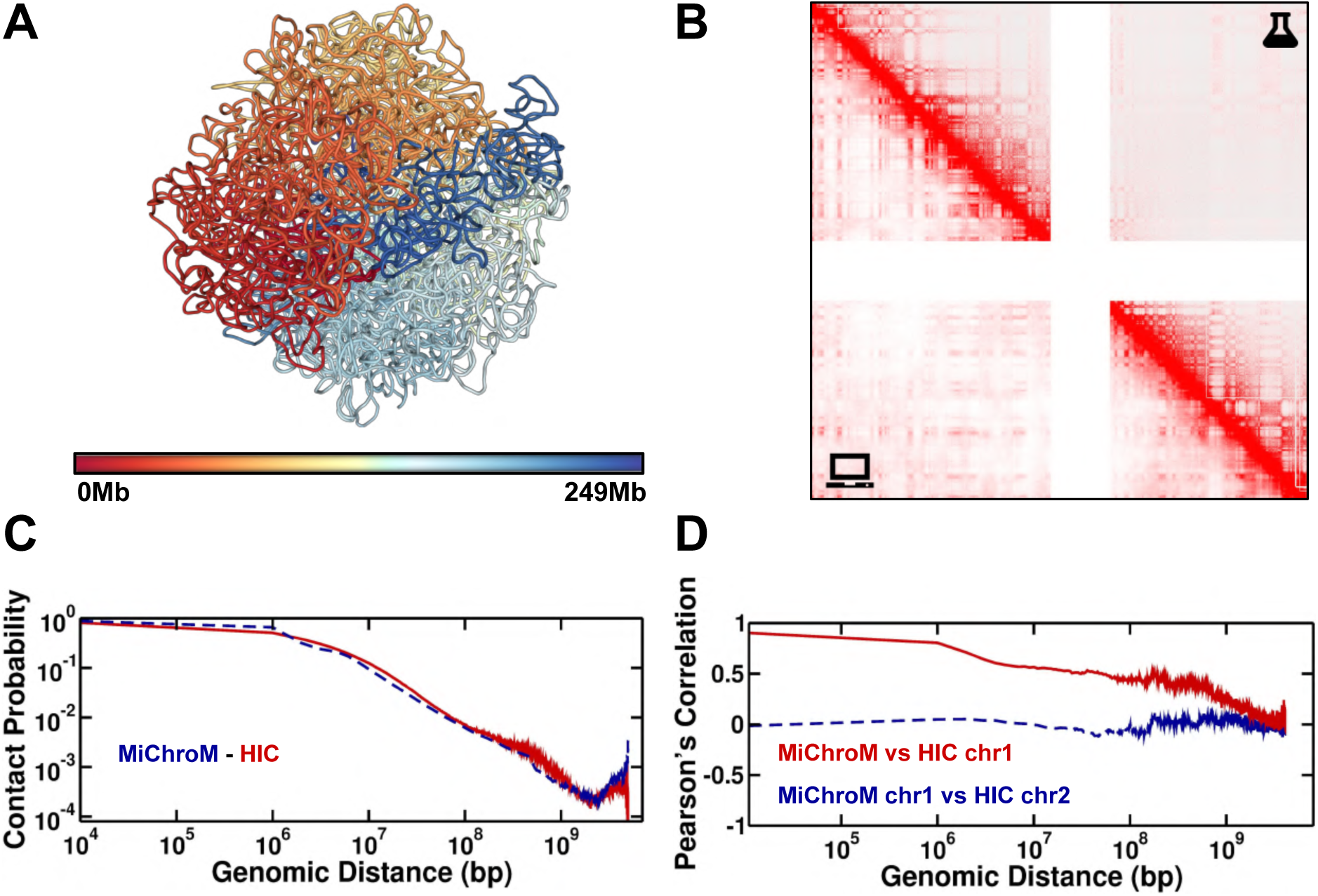
The structural ensemble of chromosome 1, cell line A549, generated by simulations using the MiChroM+MEGABASE pipeline. A - Three-dimensional representative structure colored by index, red to blue. B - Comparison between Hi-C maps obtained from wet-lab experiments (top) and *in silico* (bottom). The Hi-C maps predicted from simulations have been sampled from an ensemble of 500 thousand structures. C - Contact probability as a function of the genomic distance. The solid red line is the curve extracted from the experimental data. The dashed blue line is the data obtained from the *in silico* HiC. D - Pearson’s correlation between experimental and simulated Hi-C maps of chromosome 1 as a function of the genomic distance, shown by the solid red line. As a term of comparison the dashed blue line shows the correlation between Hi-C maps of different chromosomes.

**Figure S2:**
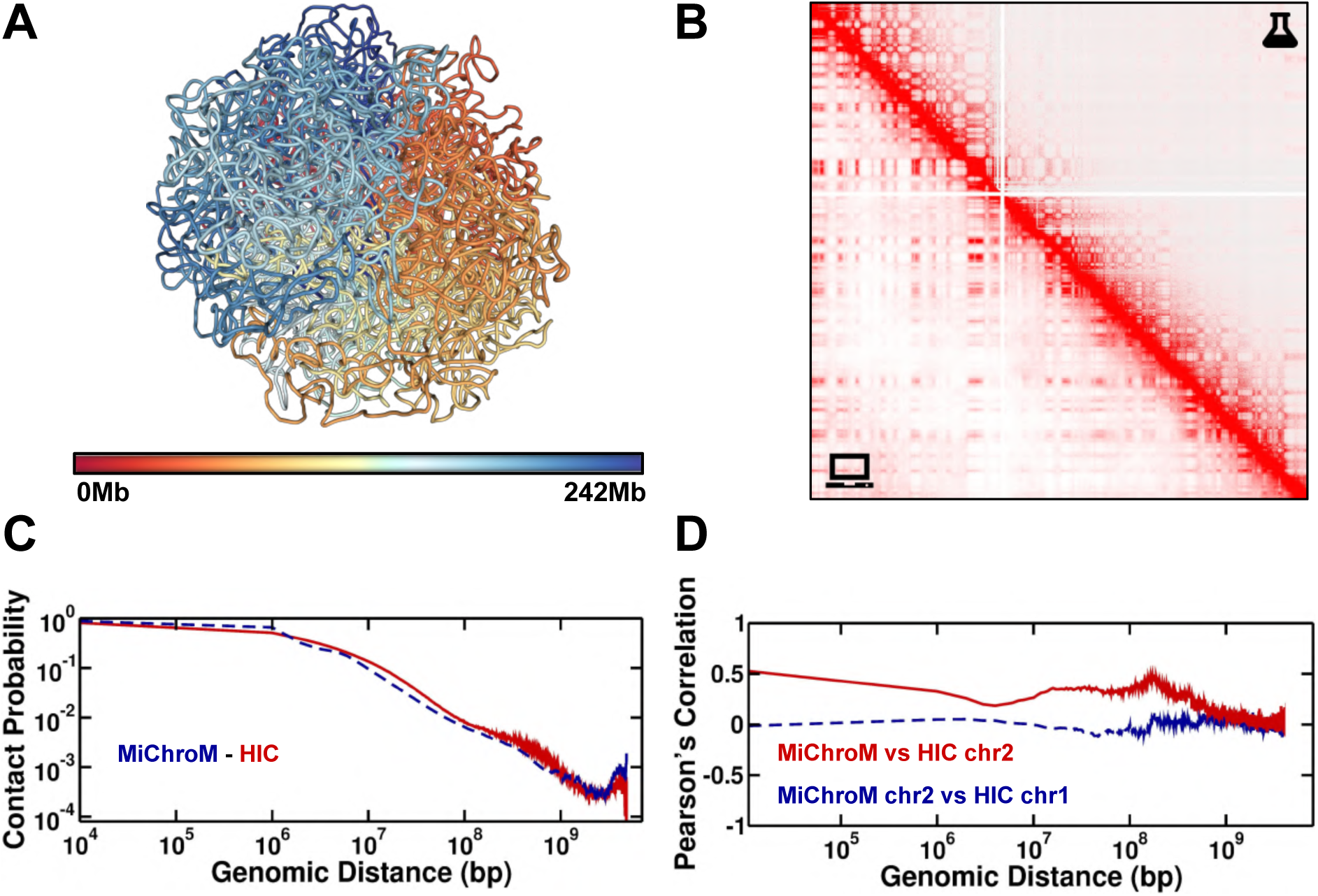
The structural ensemble of chromosome 2, cell line A549, generated by simulations using the MiChroM+MEGABASE pipeline. A - Three-dimensional representative structure colored by index, red to blue. B - Comparison between Hi-C maps obtained from wet-lab experiments (top) and *in silico* (bottom). The Hi-C maps predicted from simulations have been sampled from an ensemble of 500 thousand structures. C - Contact probability as a function of the genomic distance. The solid red line is the curve extracted from the experimental data. The dashed blue line is the data obtained from the *in silico* HiC. D - Pearson’s correlation between experimental and simulated Hi-C maps of chromosome 2 as a function of the genomic distance, shown by the solid red line. As a term of comparison the dashed blue line shows the correlation between Hi-C maps of different chromosomes.

**Figure S3:**
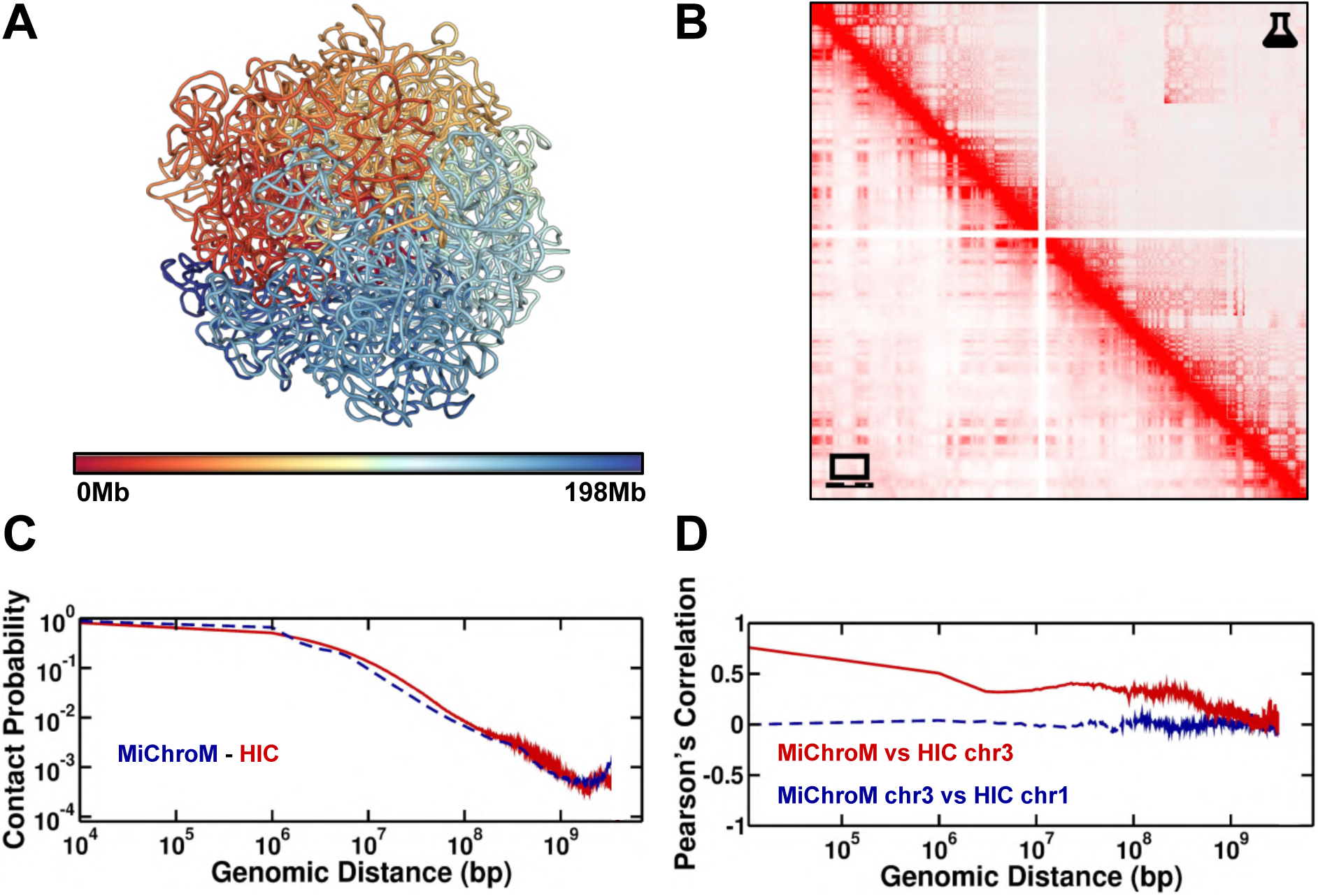
The structural ensemble of chromosome 3, cell line A549, generated by simulations using the MiChroM+MEGABASE pipeline. A - Three-dimensional representative structure colored by index, red to blue. B - Comparison between Hi-C maps obtained from wet-lab experiments (top) and *in silico* (bottom). The Hi-C maps predicted from simulations have been sampled from an ensemble of 500 thousand structures. C - Contact probability as a function of the genomic distance. The solid red line is the curve extracted from the experimental data. The dashed blue line is the data obtained from the *in silico* HiC. D - Pearson’s correlation between experimental and simulated Hi-C maps of chromosome 3 as a function of the genomic distance, shown by the solid red line. As a term of comparison the dashed blue line shows the correlation between Hi-C maps of different chromosomes.

**Figure S4:**
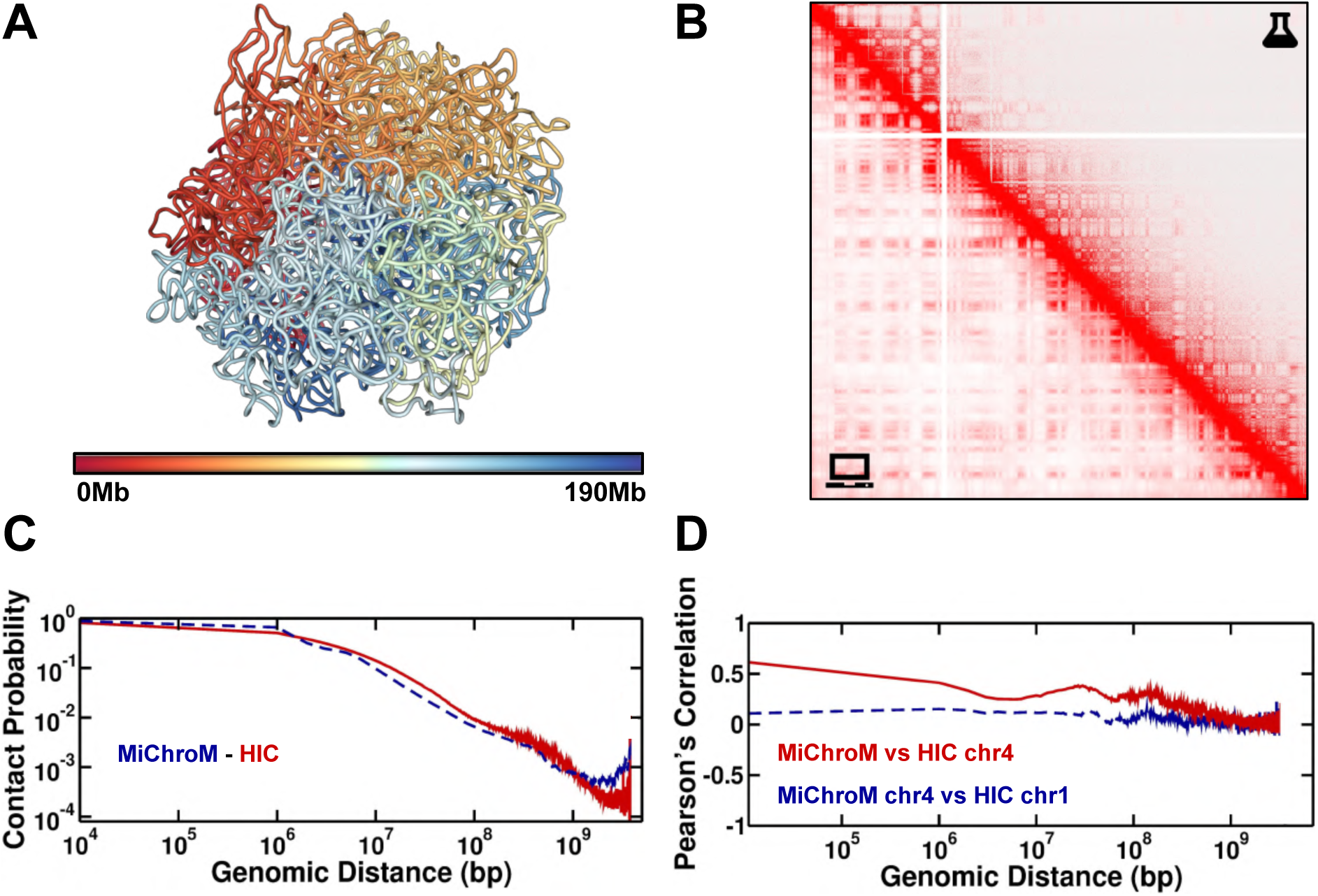
The structural ensemble of chromosome 4, cell line A549, generated by simulations using the MiChroM+MEGABASE pipeline. A - Three-dimensional representative structure colored by index, red to blue. B - Comparison between Hi-C maps obtained from wet-lab experiments (top) and *in silico* (bottom). The Hi-C maps predicted from simulations have been sampled from an ensemble of 500 thousand structures. C - Contact probability as a function of the genomic distance. The solid red line is the curve extracted from the experimental data. The dashed blue line is the data obtained from the *in silico* HiC. D - Pearson’s correlation between experimental and simulated Hi-C maps of chromosome 4 as a function of the genomic distance, shown by the solid red line. As a term of comparison the dashed blue line shows the correlation between Hi-C maps of different chromosomes.

**Figure S5:**
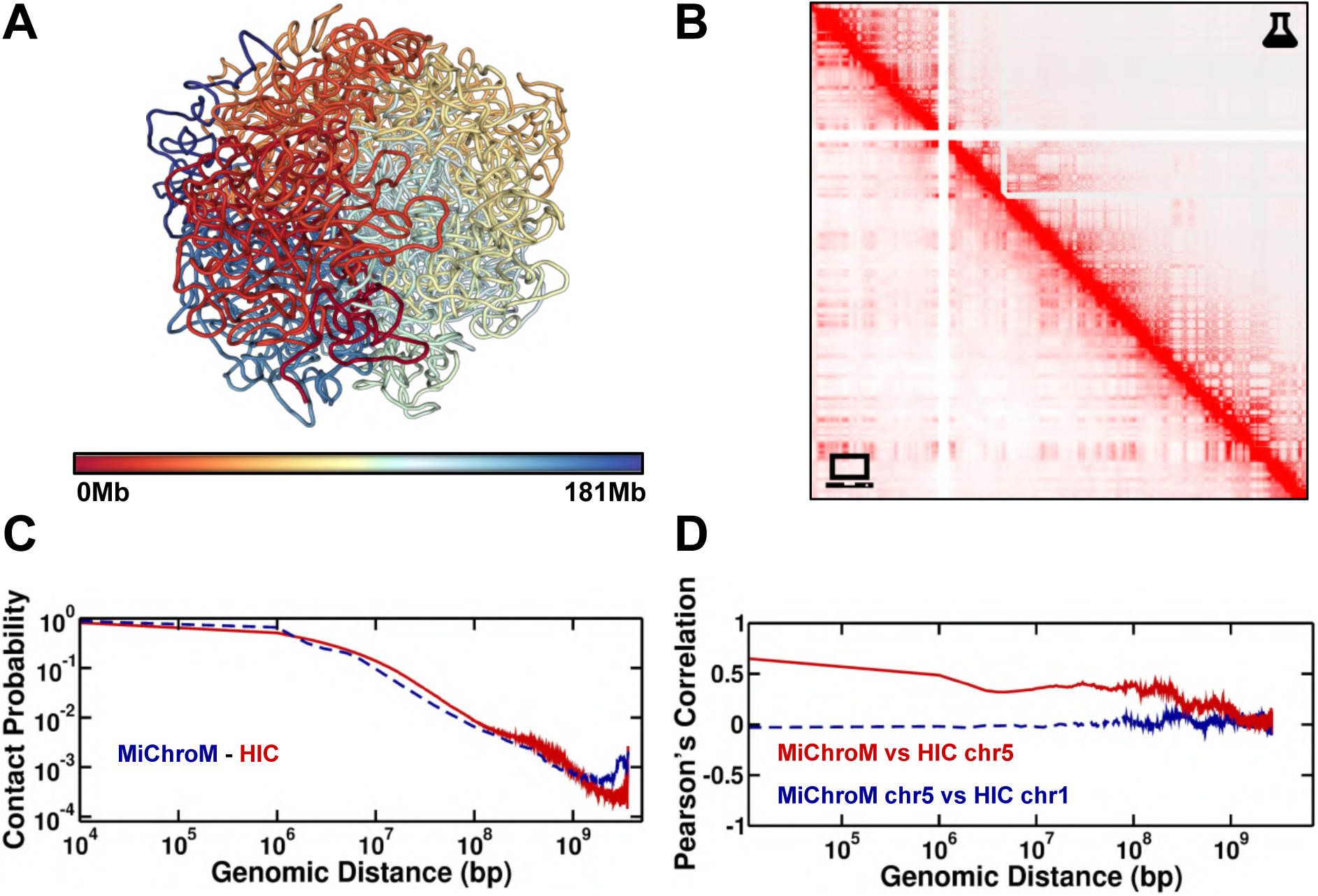
The structural ensemble of chromosome 5, cell line A549, generated by simulations using the MiChroM+MEGABASE pipeline. A - Three-dimensional representative structure colored by index, red to blue. B - Comparison between Hi-C maps obtained from wet-lab experiments (top) and *in silico* (bottom). The Hi-C maps predicted from simulations have been sampled from an ensemble of 500 thousand structures. C - Contact probability as a function of the genomic distance. The solid red line is the curve extracted from the experimental data. The dashed blue line is the data obtained from the *in silico* HiC. D - Pearson’s correlation between experimental and simulated Hi-C maps of chromosome 5 as a function of the genomic distance, shown by the solid red line. As a term of comparison the dashed blue line shows the correlation between Hi-C maps of different chromosomes.

**Figure S6:**
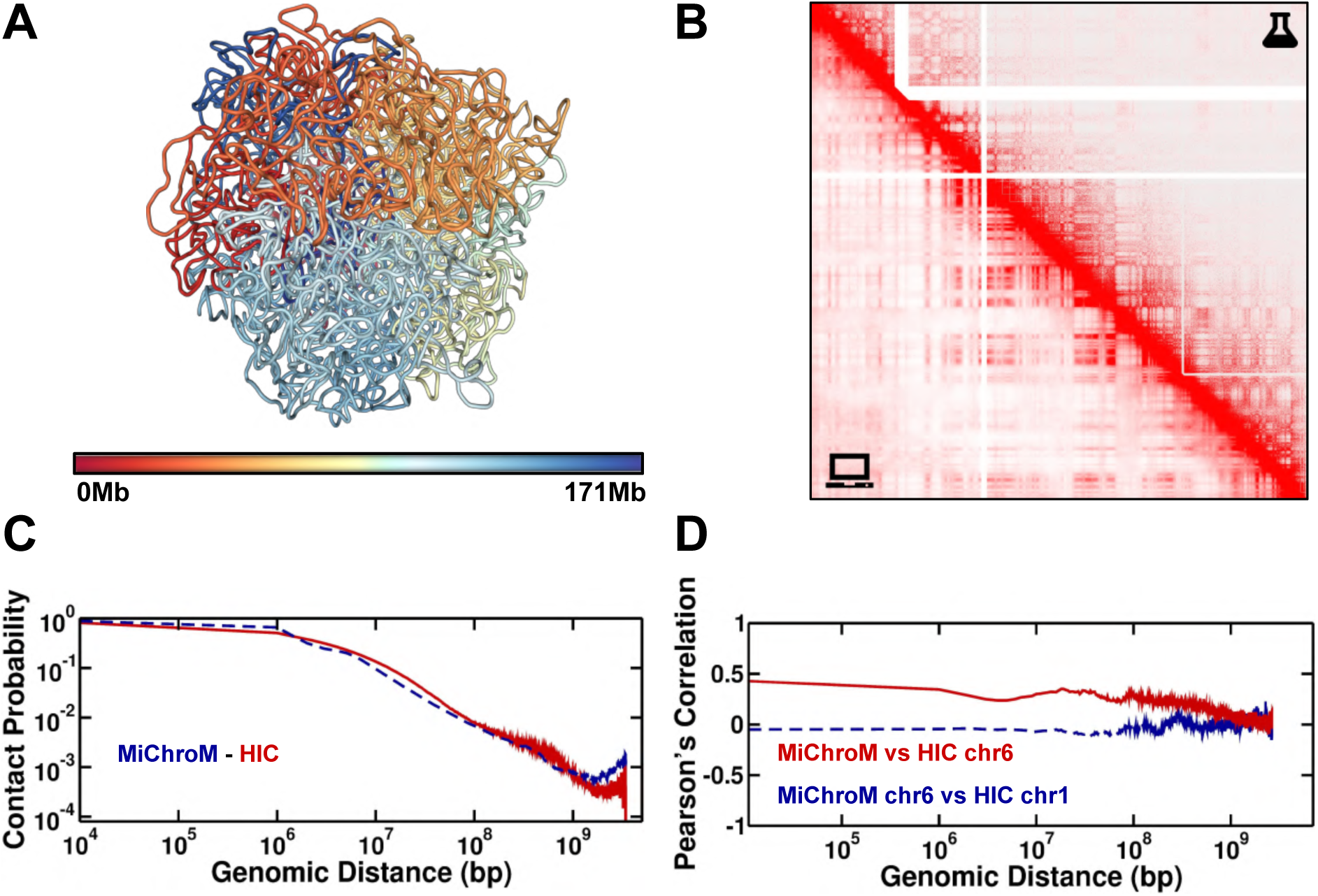
The structural ensemble of chromosome 6, cell line A549, generated by simulations using the MiChroM+MEGABASE pipeline. A - Three-dimensional representative structure colored by index, red to blue. B - Comparison between Hi-C maps obtained from wet-lab experiments (top) and *in silico* (bottom). The Hi-C maps predicted from simulations have been sampled from an ensemble of 500 thousand structures. C - Contact probability as a function of the genomic distance. The solid red line is the curve extracted from the experimental data. The dashed blue line is the data obtained from the *in silico* HiC. D - Pearson’s correlation between experimental and simulated Hi-C maps of chromosome 6 as a function of the genomic distance, shown by the solid red line. As a term of comparison the dashed blue line shows the correlation between Hi-C maps of different chromosomes.

**Figure S7:**
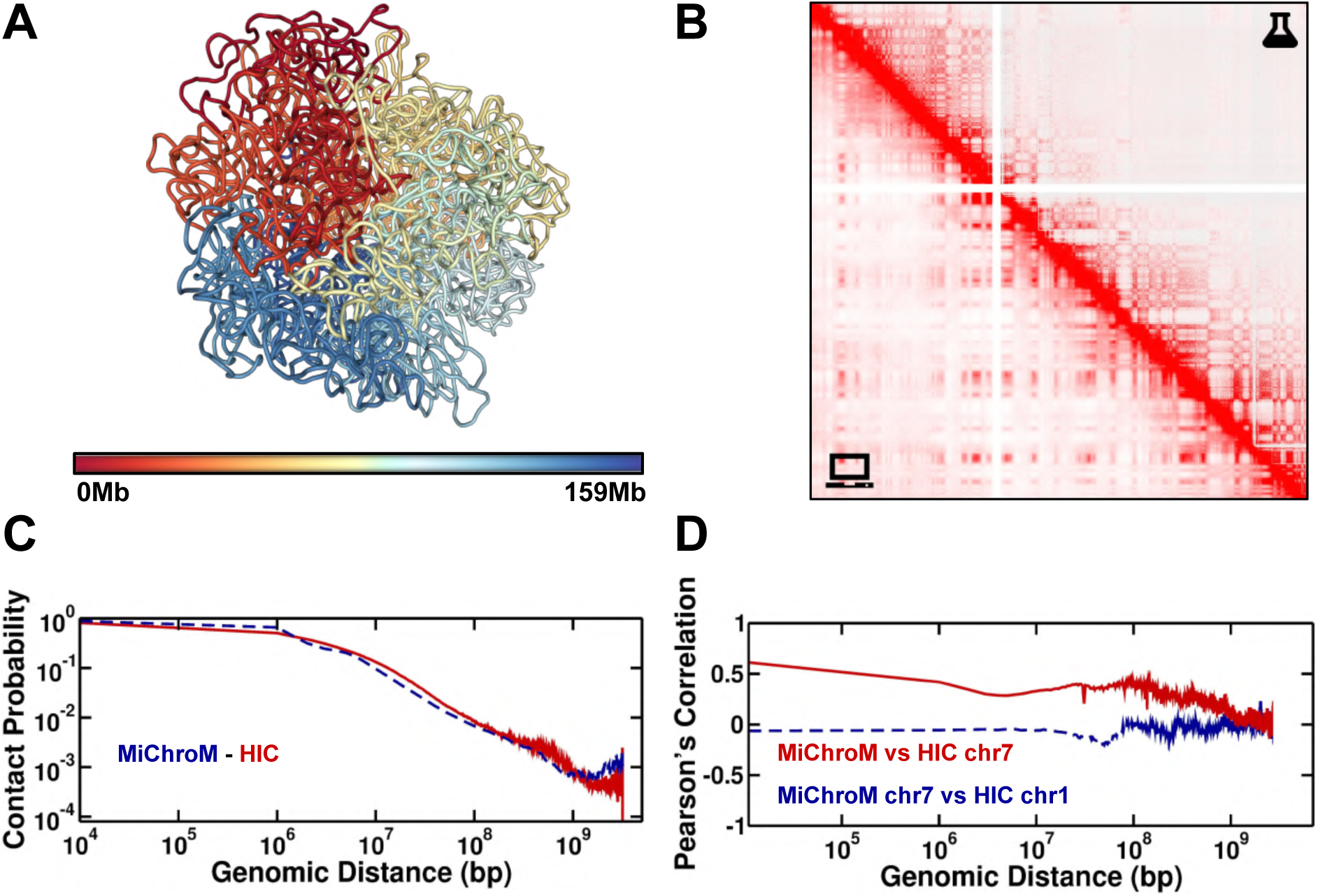
The structural ensemble of chromosome 7, cell line A549, generated by simulations using the MiChroM+MEGABASE pipeline. A - Three-dimensional representative structure colored by index, red to blue. B - Comparison between Hi-C maps obtained from wet-lab experiments (top) and *in silico* (bottom). The Hi-C maps predicted from simulations have been sampled from an ensemble of 500 thousand structures. C - Contact probability as a function of the genomic distance. The solid red line is the curve extracted from the experimental data. The dashed blue line is the data obtained from the *in silico* HiC. D - Pearson’s correlation between experimental and simulated Hi-C maps of chromosome 7 as a function of the genomic distance, shown by the solid red line. As a term of comparison the dashed blue line shows the correlation between Hi-C maps of different chromosomes.

**Figure S8:**
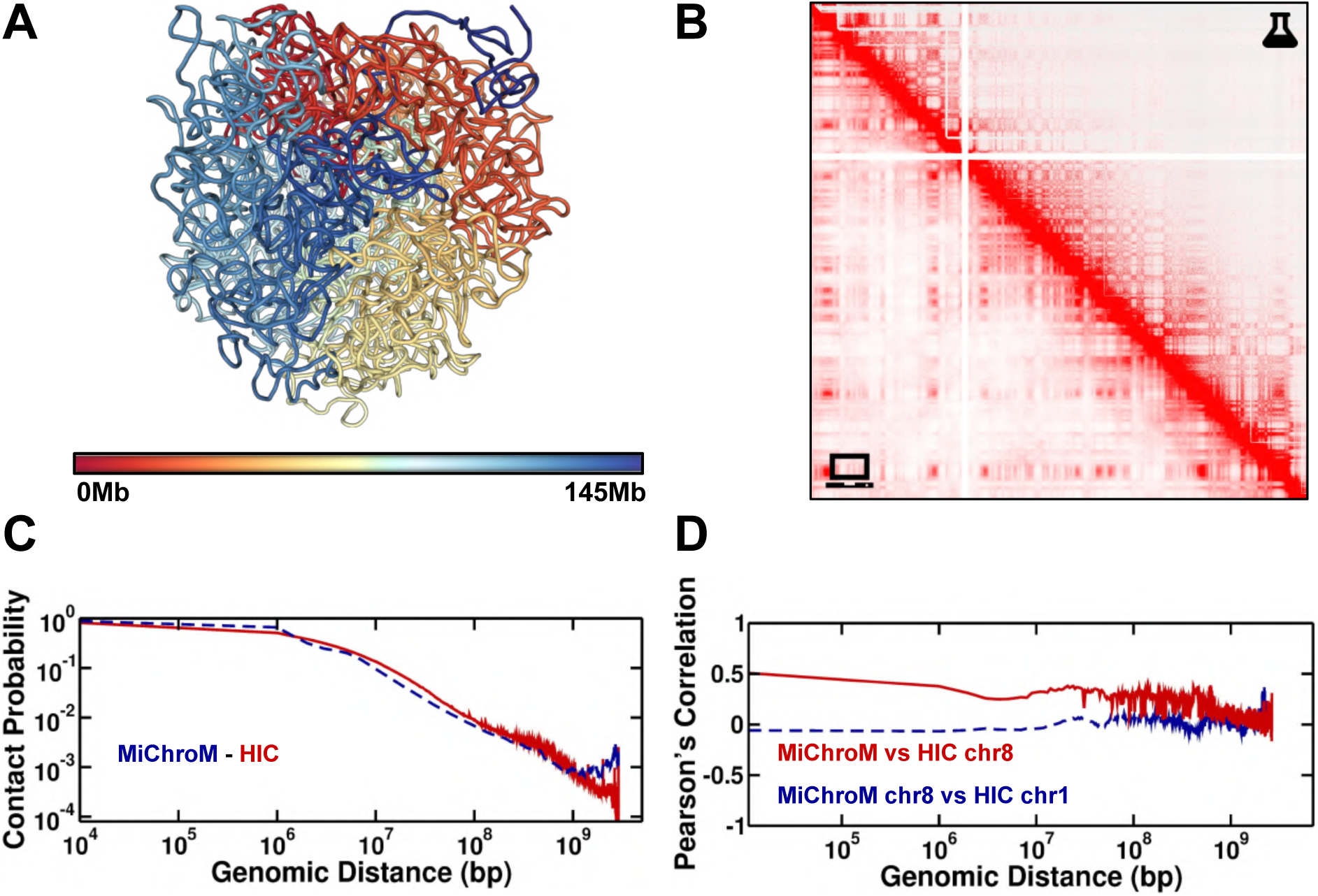
The structural ensemble of chromosome 8, cell line A549, generated by simulations using the MiChroM+MEGABASE pipeline. A - Three-dimensional representative structure colored by index, red to blue. B - Comparison between Hi-C maps obtained from wet-lab experiments (top) and *in silico* (bottom). The Hi-C maps predicted from simulations have been sampled from an ensemble of 500 thousand structures. C - Contact probability as a function of the genomic distance. The solid red line is the curve extracted from the experimental data. The dashed blue line is the data obtained from the *in silico* HiC. D - Pearson’s correlation between experimental and simulated Hi-C maps of chromosome 8 as a function of the genomic distance, shown by the solid red line. As a term of comparison the dashed blue line shows the correlation between Hi-C maps of different chromosomes.

**Figure S9:**
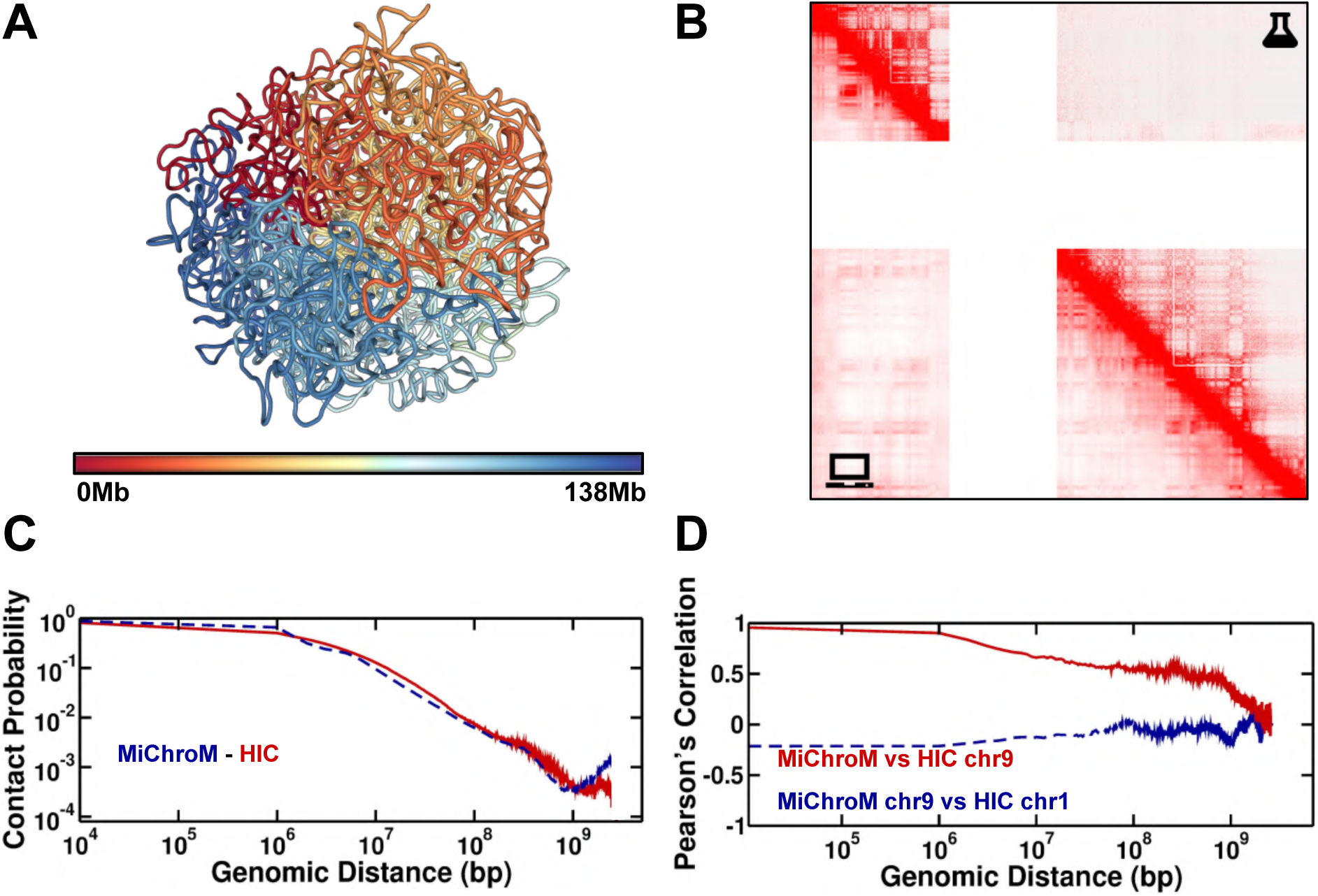
The structural ensemble of chromosome 9, cell line A549, generated by simulations using the MiChroM+MEGABASE pipeline. A - Three-dimensional representative structure colored by index, red to blue. B - Comparison between Hi-C maps obtained from wet-lab experiments (top) and *in silico* (bottom). The Hi-C maps predicted from simulations have been sampled from an ensemble of 500 thousand structures. C - Contact probability as a function of the genomic distance. The solid red line is the curve extracted from the experimental data. The dashed blue line is the data obtained from the *in silico* HiC. D - Pearson’s correlation between experimental and simulated Hi-C maps of chromosome 9 as a function of the genomic distance, shown by the solid red line. As a term of comparison the dashed blue line shows the correlation between Hi-C maps of different chromosomes.

**Figure S10:**
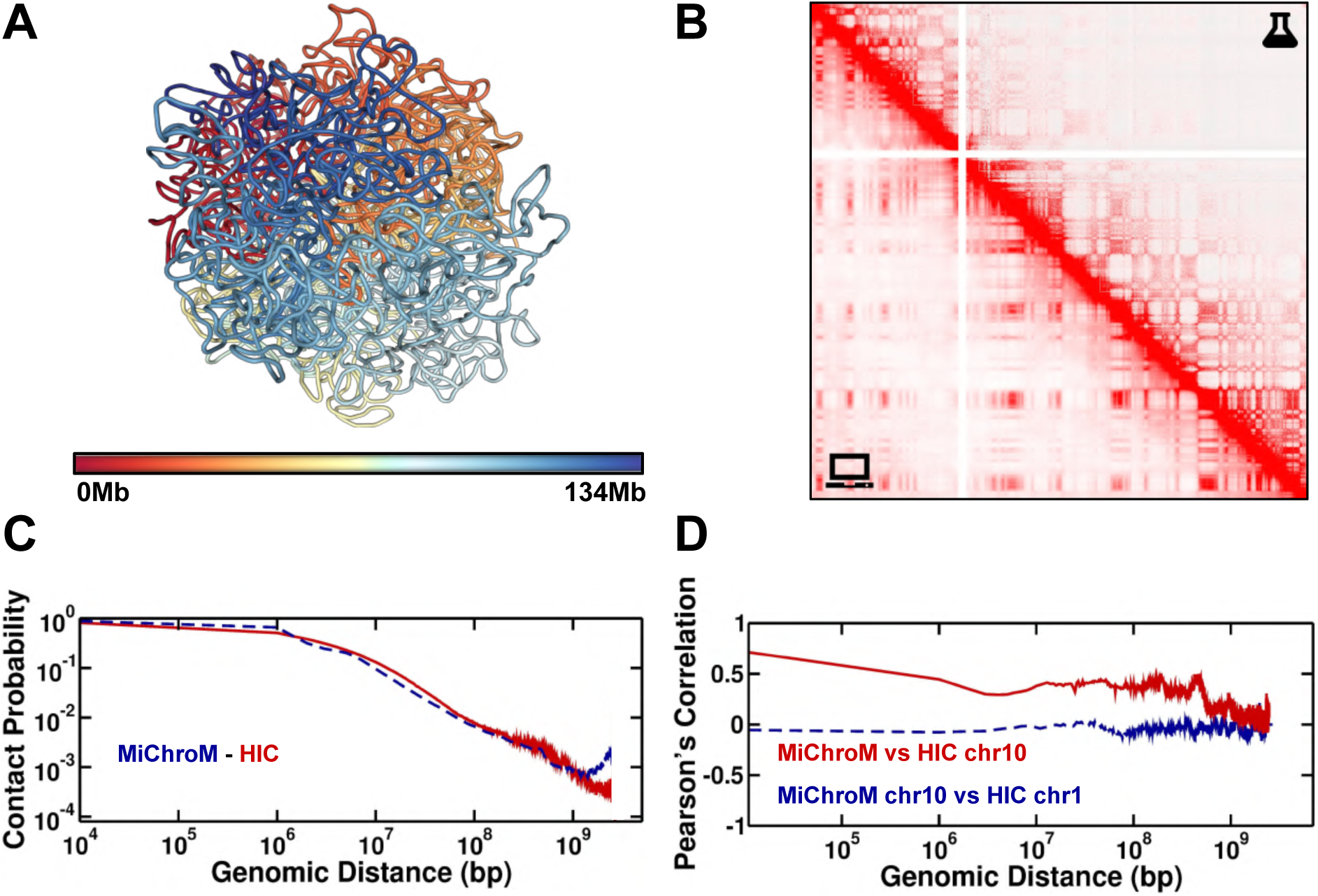
The structural ensemble of chromosome 10, cell line A549, generated by simulations using the MiChroM+MEGABASE pipeline. A - Three-dimensional representative structure colored by index, red to blue. B - Comparison between Hi-C maps obtained from wet-lab experiments (top) and *in silico* (bottom). The Hi-C maps predicted from simulations have been sampled from an ensemble of 500 thousand structures. C - Contact probability as a function of the genomic distance. The solid red line is the curve extracted from the experimental data. The dashed blue line is the data obtained from the *in silico* HiC. D - Pearson’s correlation between experimental and simulated Hi-C maps of chromosome 10 as a function of the genomic distance, shown by the solid red line. As a term of comparison the dashed blue line shows the correlation between Hi-C maps of different chromosomes.

**Figure S11:**
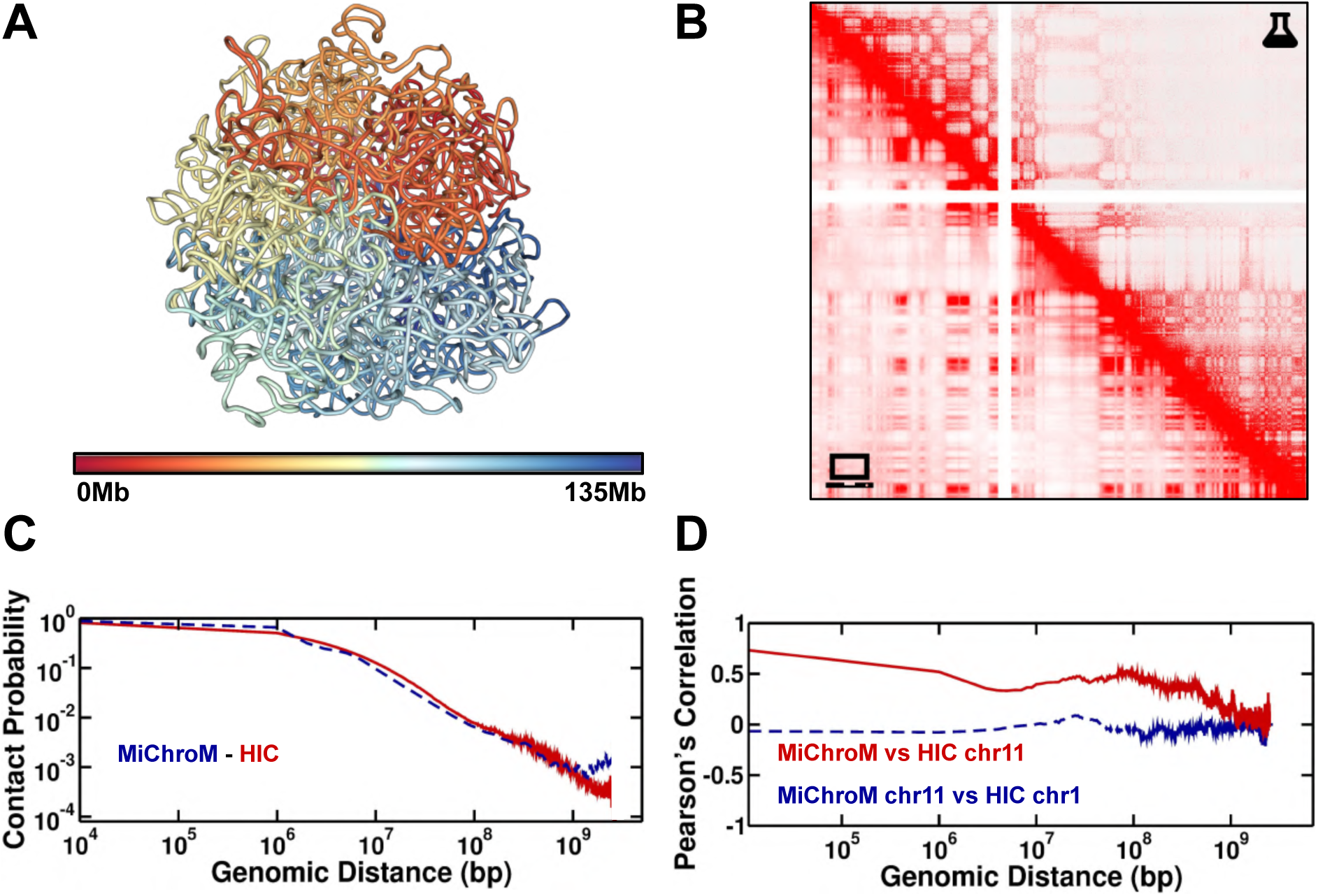
The structural ensemble of chromosome 11, cell line A549, generated by simulations using the MiChroM+MEGABASE pipeline. A - Three-dimensional representative structure colored by index, red to blue. B - Comparison between Hi-C maps obtained from wet-lab experiments (top) and *in silico* (bottom). The Hi-C maps predicted from simulations have been sampled from an ensemble of 500 thousand structures. C - Contact probability as a function of the genomic distance. The solid red line is the curve extracted from the experimental data. The dashed blue line is the data obtained from the *in silico* HiC. D - Pearson’s correlation between experimental and simulated Hi-C maps of chromosome 11 as a function of the genomic distance, shown by the solid red line. As a term of comparison the dashed blue line shows the correlation between Hi-C maps of different chromosomes.

**Figure S12:**
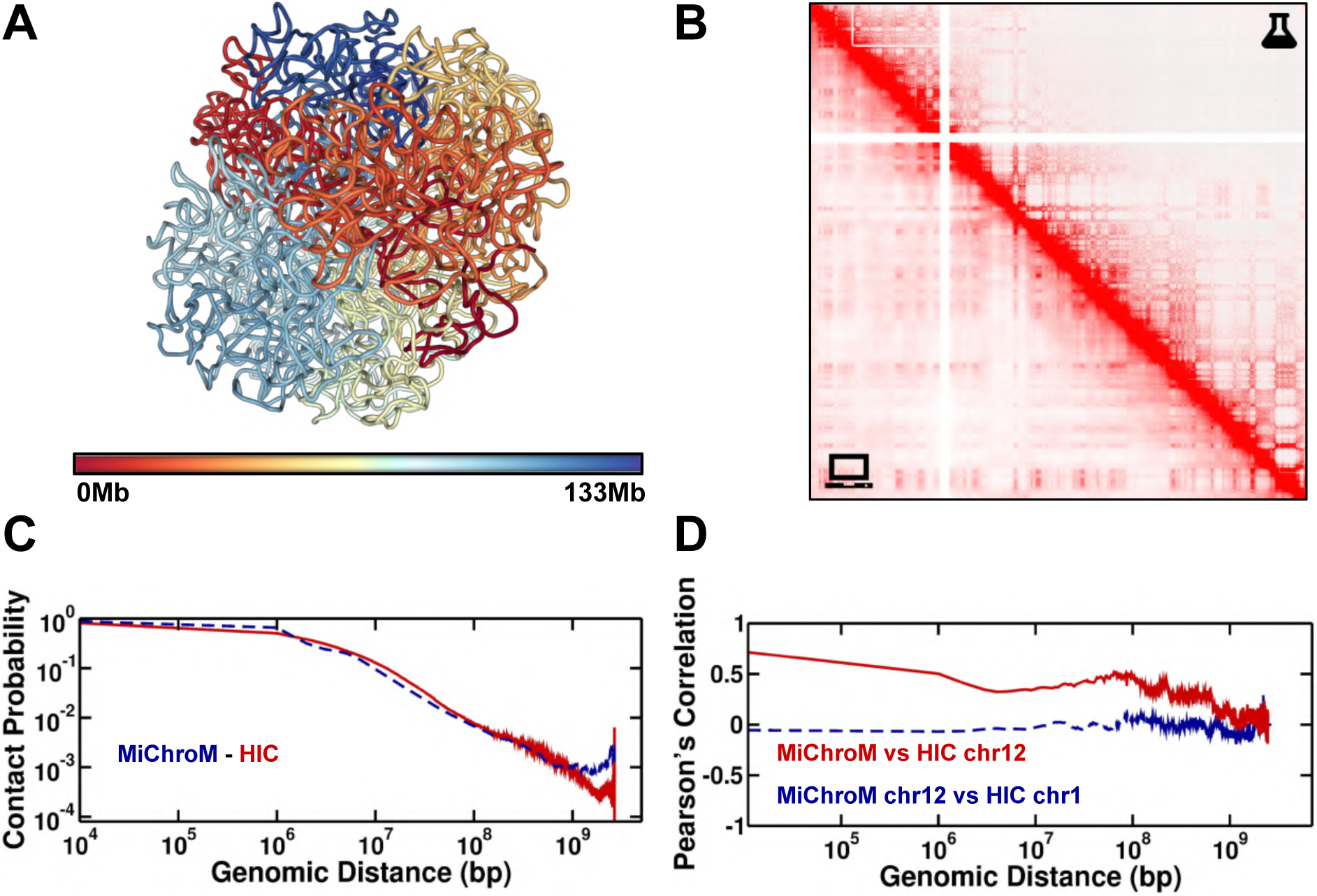
The structural ensemble of chromosome 12, cell line A549, generated by simulations using the MiChroM+MEGABASE pipeline. A - Three-dimensional representative structure colored by index, red to blue. B - Comparison between Hi-C maps obtained from wet-lab experiments (top) and *in silico* (bottom). The Hi-C maps predicted from simulations have been sampled from an ensemble of 500 thousand structures. C - Contact probability as a function of the genomic distance. The solid red line is the curve extracted from the experimental data. The dashed blue line is the data obtained from the *in silico* HiC. D - Pearson’s correlation between experimental and simulated Hi-C maps of chromosome 12 as a function of the genomic distance, shown by the solid red line. As a term of comparison the dashed blue line shows the correlation between Hi-C maps of different chromosomes.

**Figure S13:**
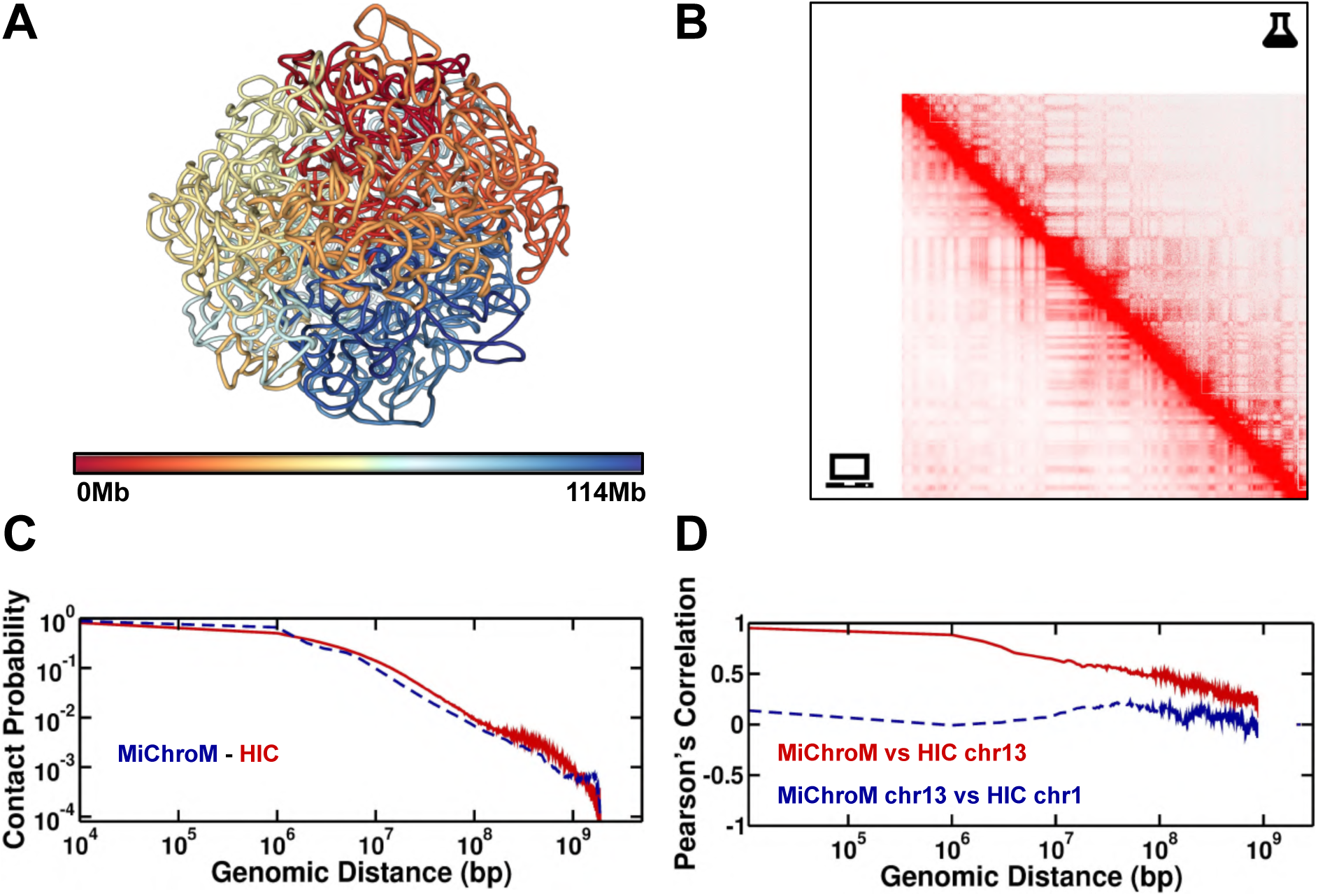
The structural ensemble of chromosome 13, cell line A549, generated by simulations using the MiChroM+MEGABASE pipeline. A - Three-dimensional representative structure colored by index, red to blue. B - Comparison between Hi-C maps obtained from wet-lab experiments (top) and *in silico* (bottom). The Hi-C maps predicted from simulations have been sampled from an ensemble of 500 thousand structures. C - Contact probability as a function of the genomic distance. The solid red line is the curve extracted from the experimental data. The dashed blue line is the data obtained from the *in silico* HiC. D - Pearson’s correlation between experimental and simulated Hi-C maps of chromosome 13 as a function of the genomic distance, shown by the solid red line. As a term of comparison the dashed blue line shows the correlation between Hi-C maps of different chromosomes.

**Figure S14:**
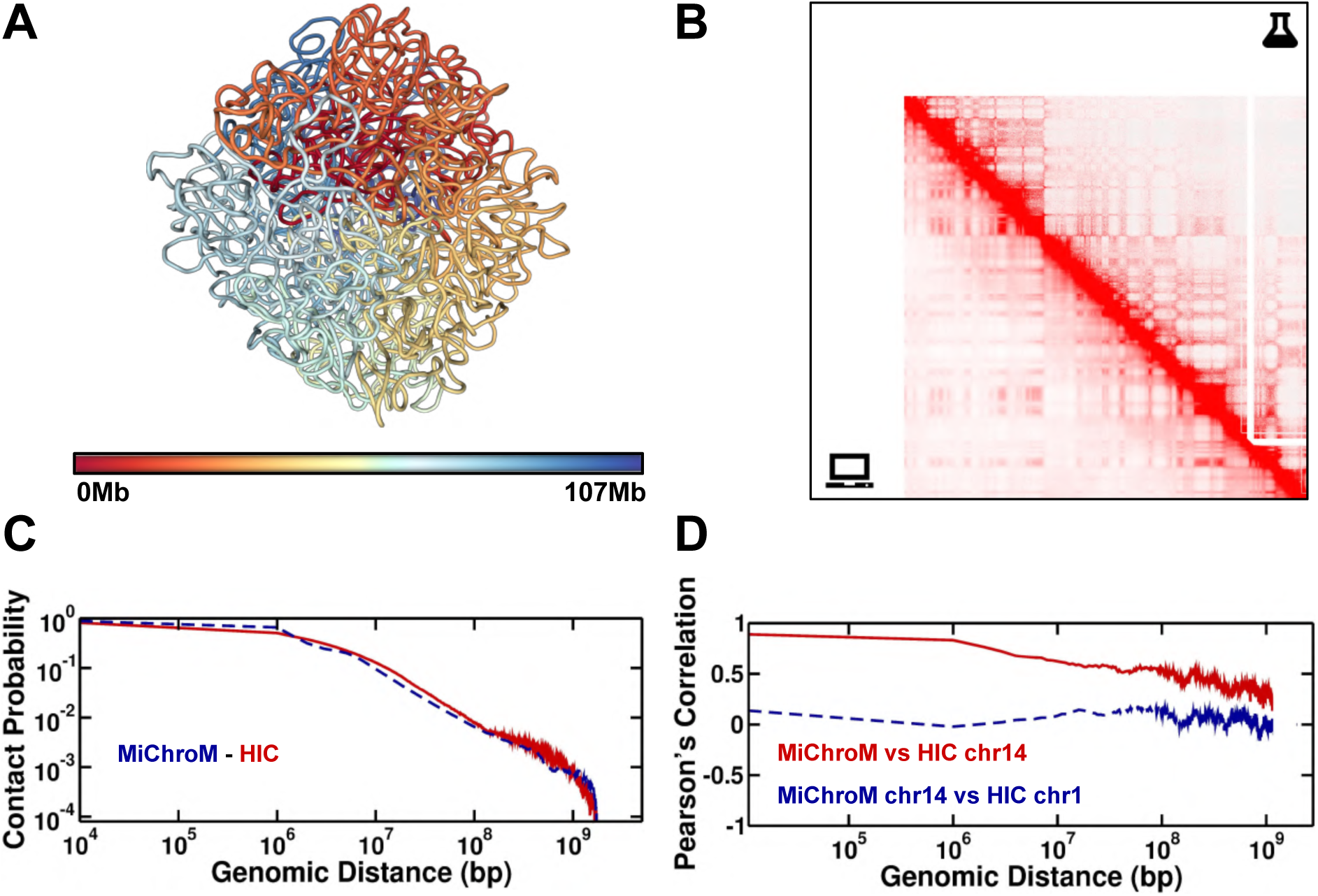
The structural ensemble of chromosome 14, cell line A549, generated by simulations using the MiChroM+MEGABASE pipeline. A - Three-dimensional representative structure colored by index, red to blue. B - Comparison between Hi-C maps obtained from wet-lab experiments (top) and *in silico* (bottom). The Hi-C maps predicted from simulations have been sampled from an ensemble of 500 thousand structures. C - Contact probability as a function of the genomic distance. The solid red line is the curve extracted from the experimental data. The dashed blue line is the data obtained from the *in silico* HiC. D - Pearson’s correlation between experimental and simulated Hi-C maps of chromosome 14 as a function of the genomic distance, shown by the solid red line. As a term of comparison the dashed blue line shows the correlation between Hi-C maps of different chromosomes.

**Figure S15:**
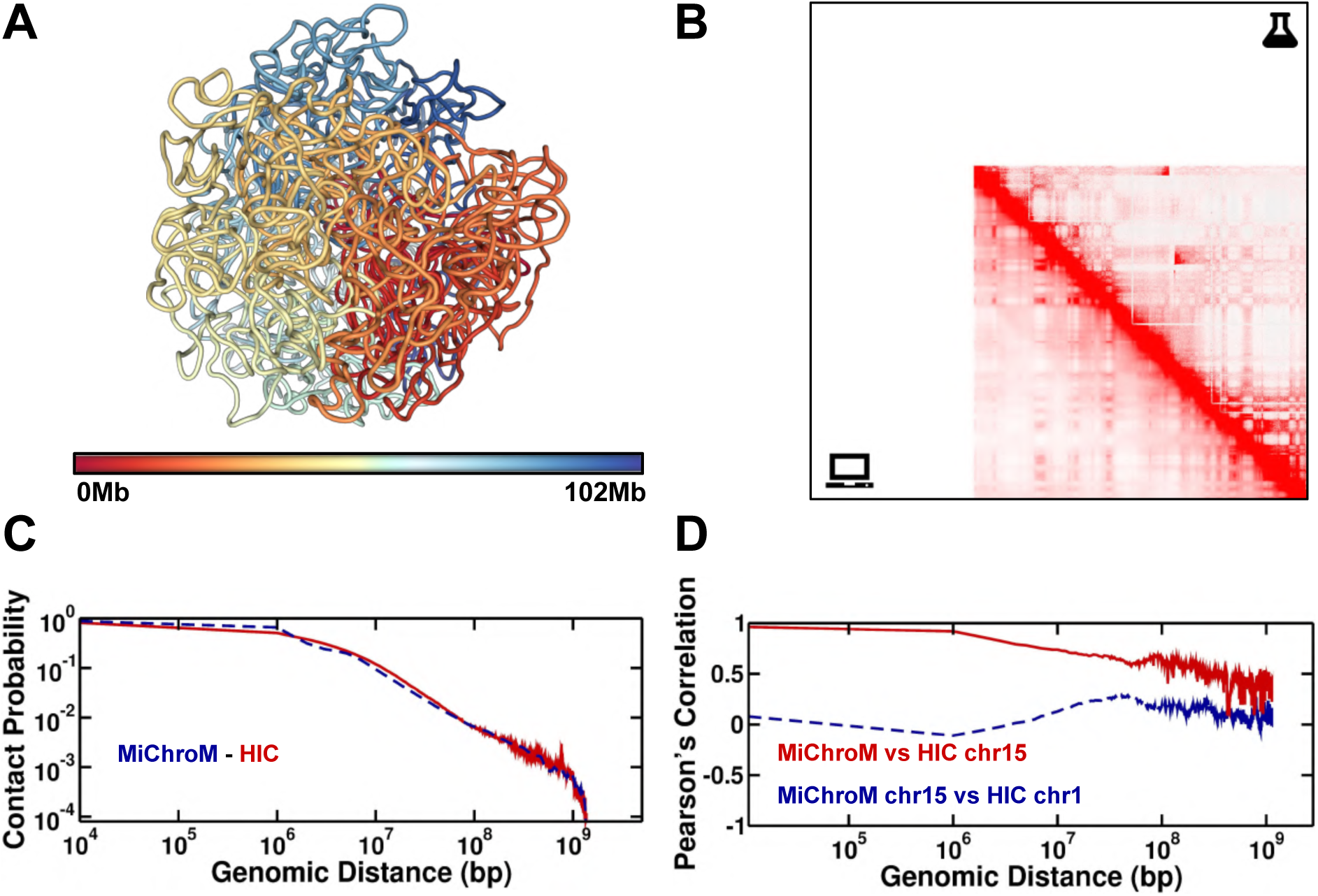
The structural ensemble of chromosome 15, cell line A549, generated by simulations using the MiChroM+MEGABASE pipeline. A - Three-dimensional representative structure colored by index, red to blue. B - Comparison between Hi-C maps obtained from wet-lab experiments (top) and *in silico* (bottom). The Hi-C maps predicted from simulations have been sampled from an ensemble of 500 thousand structures. C - Contact probability as a function of the genomic distance. The solid red line is the curve extracted from the experimental data. The dashed blue line is the data obtained from the *in silico* HiC. D - Pearson’s correlation between experimental and simulated Hi-C maps of chromosome 15 as a function of the genomic distance, shown by the solid red line. As a term of comparison the dashed blue line shows the correlation between Hi-C maps of different chromosomes.

**Figure S16:**
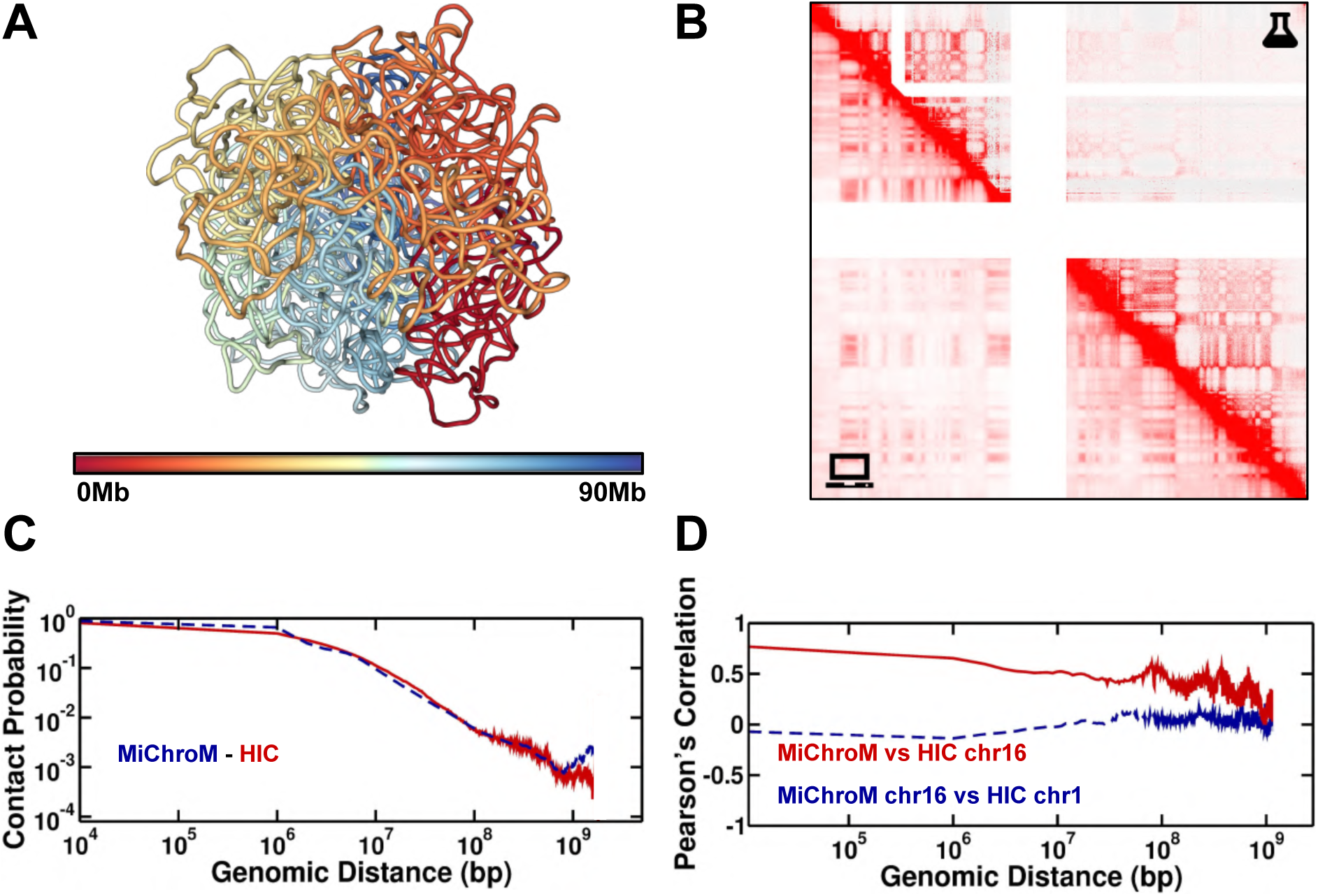
The structural ensemble of chromosome 16, cell line A549, generated by simulations using the MiChroM+MEGABASE pipeline. A - Three-dimensional representative structure colored by index, red to blue. B - Comparison between Hi-C maps obtained from wet-lab experiments (top) and *in silico* (bottom). The Hi-C maps predicted from simulations have been sampled from an ensemble of 500 thousand structures. C - Contact probability as a function of the genomic distance. The solid red line is the curve extracted from the experimental data. The dashed blue line is the data obtained from the *in silico* HiC. D - Pearson’s correlation between experimental and simulated Hi-C maps of chromosome 16 as a function of the genomic distance, shown by the solid red line. As a term of comparison the dashed blue line shows the correlation between Hi-C maps of different chromosomes.

**Figure S17:**
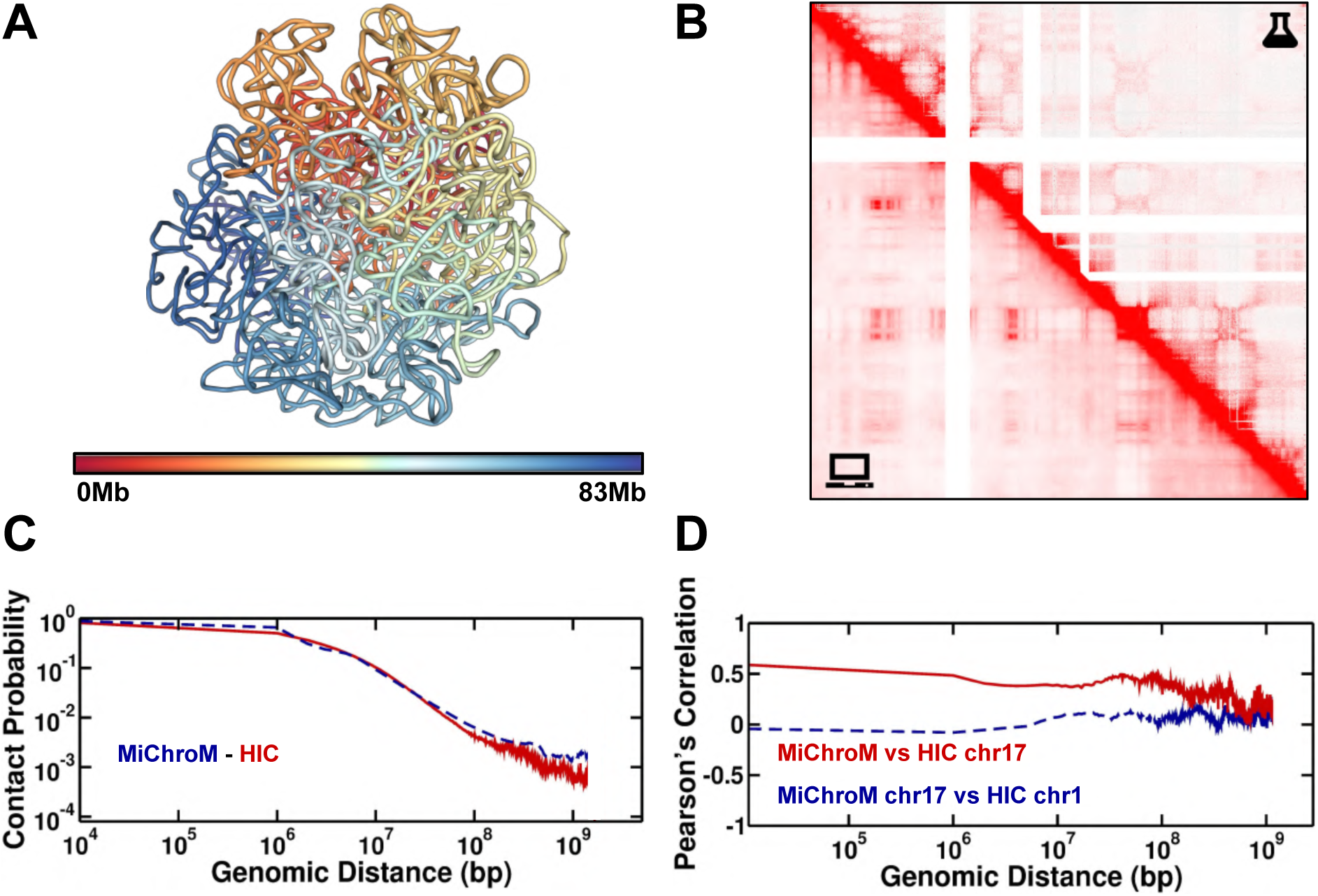
The structural ensemble of chromosome 17, cell line A549, generated by simulations using the MiChroM+MEGABASE pipeline. A - Three-dimensional representative structure colored by index, red to blue. B - Comparison between Hi-C maps obtained from wet-lab experiments (top) and *in silico* (bottom). The Hi-C maps predicted from simulations have been sampled from an ensemble of 500 thousand structures. C - Contact probability as a function of the genomic distance. The solid red line is the curve extracted from the experimental data. The dashed blue line is the data obtained from the *in silico* HiC. D - Pearson’s correlation between experimental and simulated Hi-C maps of chromosome 17 as a function of the genomic distance, shown by the solid red line. As a term of comparison the dashed blue line shows the correlation between Hi-C maps of different chromosomes.

**Figure S18:**
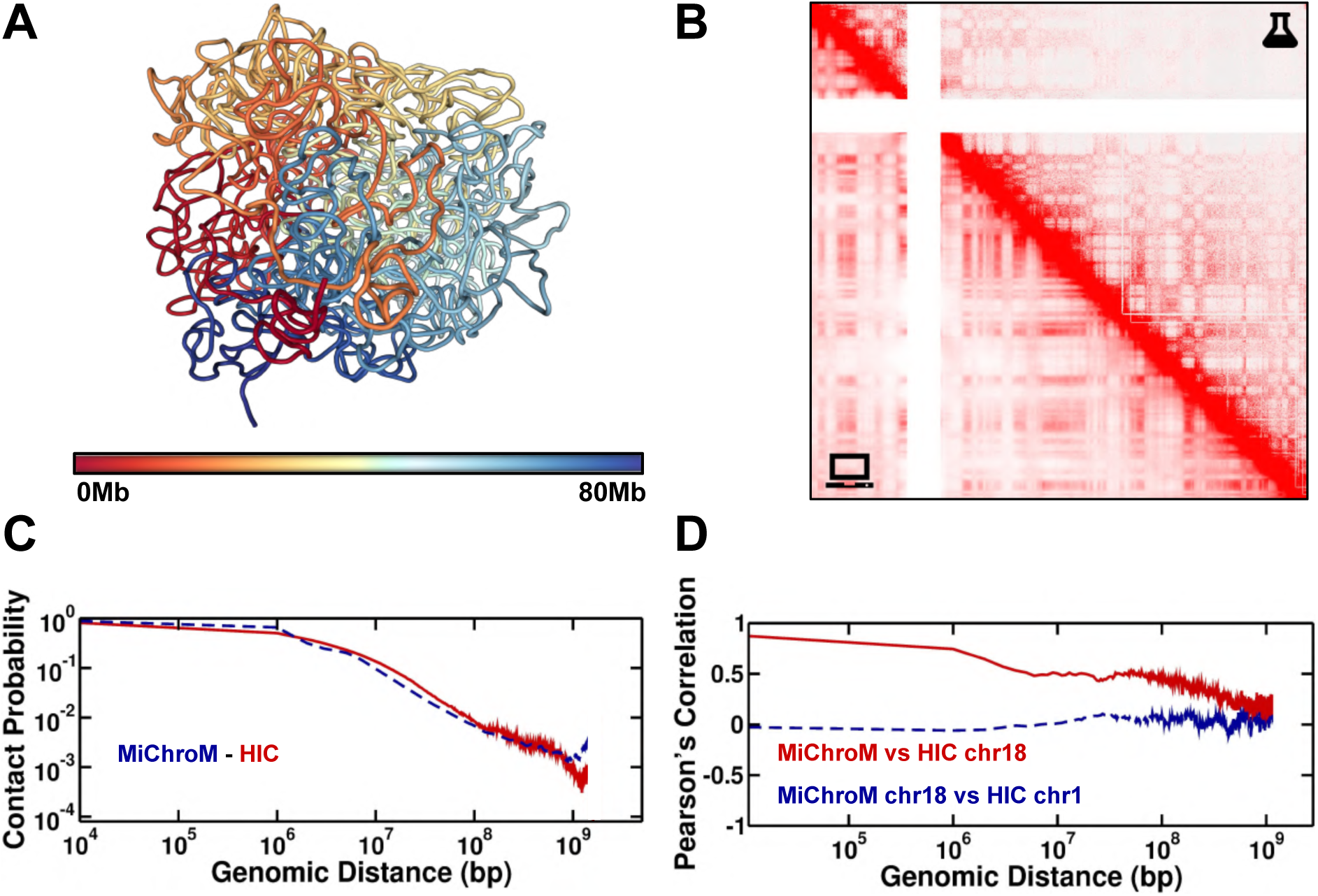
The structural ensemble of chromosome 18, cell line A549, generated by simulations using the MiChroM+MEGABASE pipeline. A - Three-dimensional representative structure colored by index, red to blue. B - Comparison between Hi-C maps obtained from wet-lab experiments (top) and *in silico* (bottom). The Hi-C maps predicted from simulations have been sampled from an ensemble of 500 thousand structures. C - Contact probability as a function of the genomic distance. The solid red line is the curve extracted from the experimental data. The dashed blue line is the data obtained from the *in silico* HiC. D - Pearson’s correlation between experimental and simulated Hi-C maps of chromosome 18 as a function of the genomic distance, shown by the solid red line. As a term of comparison the dashed blue line shows the correlation between Hi-C maps of different chromosomes.

**Figure S19:**
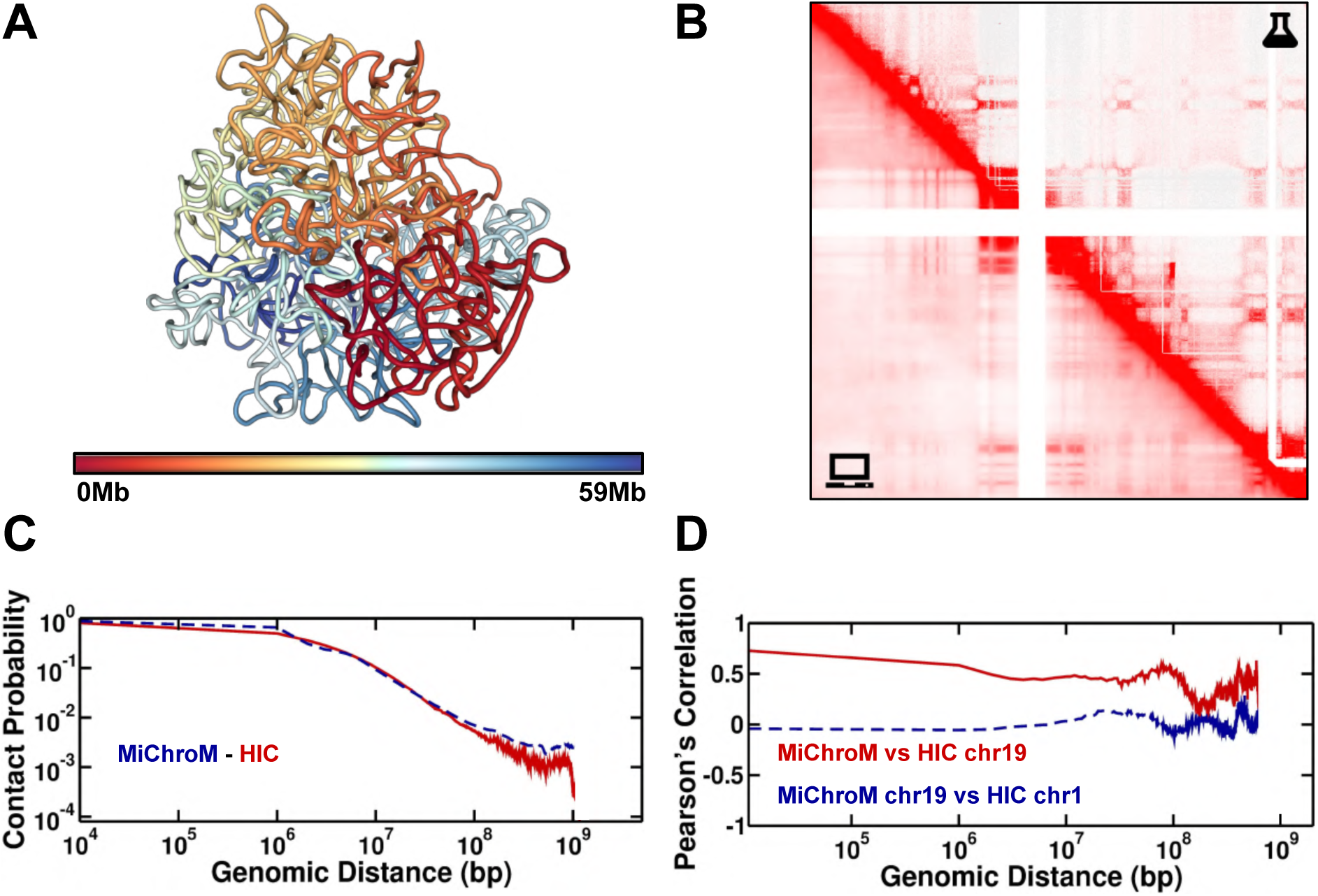
The structural ensemble of chromosome 19, cell line A549, generated by simulations using the MiChroM+MEGABASE pipeline. A - Three-dimensional representative structure colored by index, red to blue. B - Comparison between Hi-C maps obtained from wet-lab experiments (top) and *in silico* (bottom). The Hi-C maps predicted from simulations have been sampled from an ensemble of 500 thousand structures. C - Contact probability as a function of the genomic distance. The solid red line is the curve extracted from the experimental data. The dashed blue line is the data obtained from the *in silico* HiC. D - Pearson’s correlation between experimental and simulated Hi-C maps of chromosome 19 as a function of the genomic distance, shown by the solid red line. As a term of comparison the dashed blue line shows the correlation between Hi-C maps of different chromosomes.

**Figure S20:**
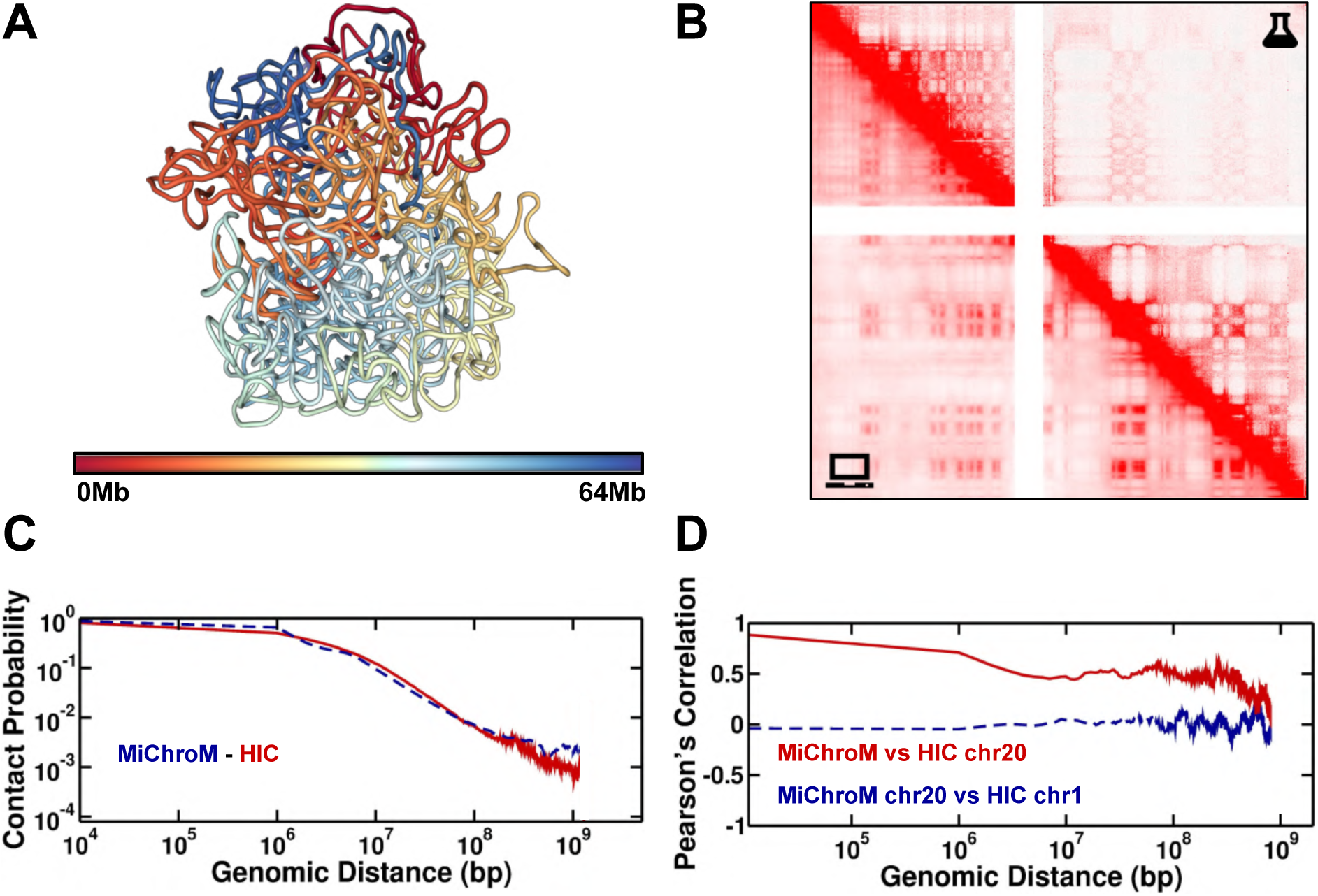
The structural ensemble of chromosome 20, cell line A549, generated by simulations using the MiChroM+MEGABASE pipeline. A - Three-dimensional representative structure colored by index, red to blue. B - Comparison between Hi-C maps obtained from wet-lab experiments (top) and *in silico* (bottom). The Hi-C maps predicted from simulations have been sampled from an ensemble of 500 thousand structures. C - Contact probability as a function of the genomic distance. The solid red line is the curve extracted from the experimental data. The dashed blue line is the data obtained from the *in silico* HiC. D - Pearson’s correlation between experimental and simulated Hi-C maps of chromosome 20 as a function of the genomic distance, shown by the solid red line. As a term of comparison the dashed blue line shows the correlation between Hi-C maps of different chromosomes.

**Figure S21:**
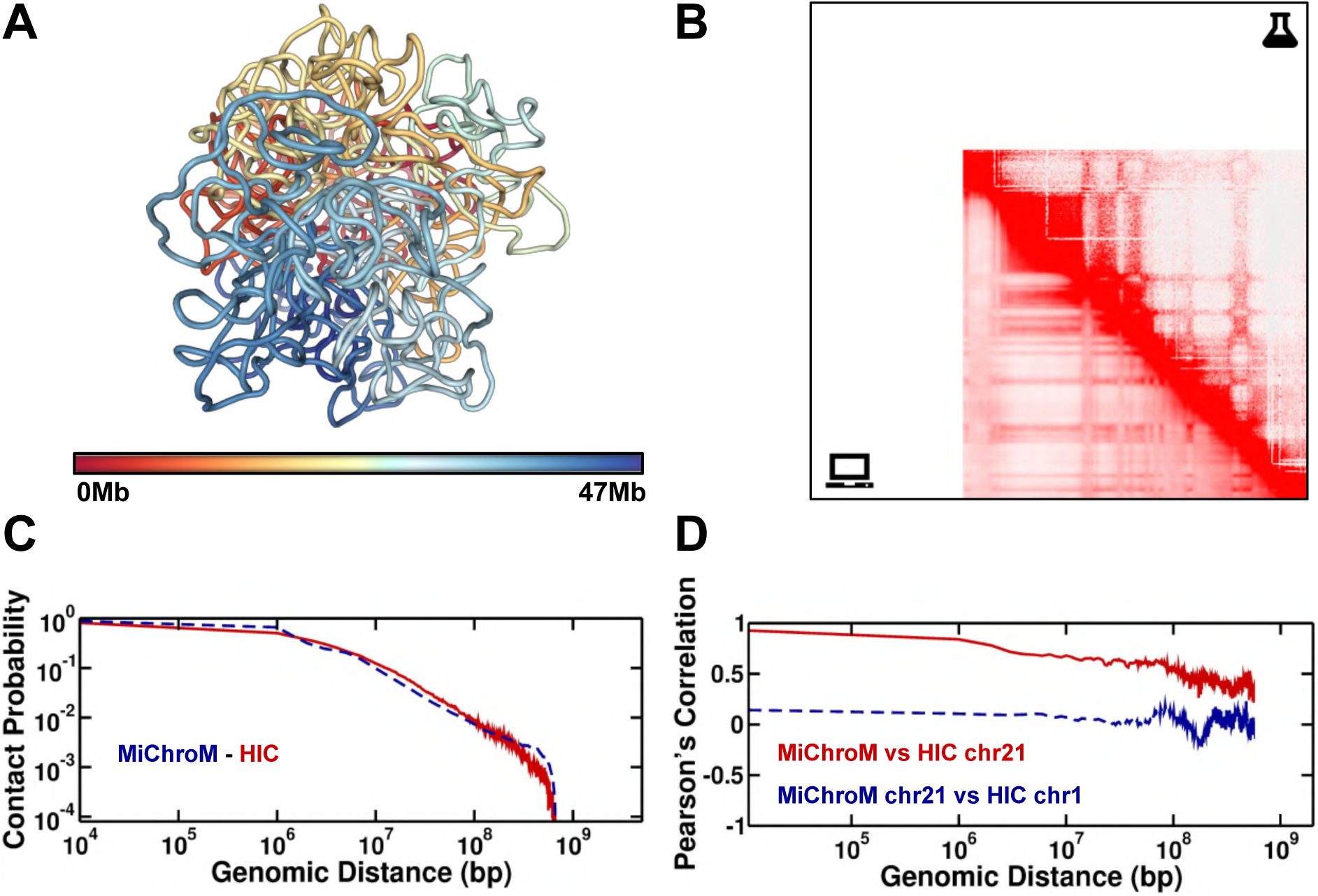
The structural ensemble of chromosome 21, cell line A549, generated by simulations using the MiChroM+MEGABASE pipeline. A - Three-dimensional representative structure colored by index, red to blue. B - Comparison between Hi-C maps obtained from wet-lab experiments (top) and *in silico* (bottom). The Hi-C maps predicted from simulations have been sampled from an ensemble of 500 thousand structures. C - Contact probability as a function of the genomic distance. The solid red line is the curve extracted from the experimental data. The dashed blue line is the data obtained from the *in silico* HiC. D - Pearson’s correlation between experimental and simulated Hi-C maps of chromosome 21 as a function of the genomic distance, shown by the solid red line. As a term of comparison the dashed blue line shows the correlation between Hi-C maps of different chromosomes.

**Figure S22:**
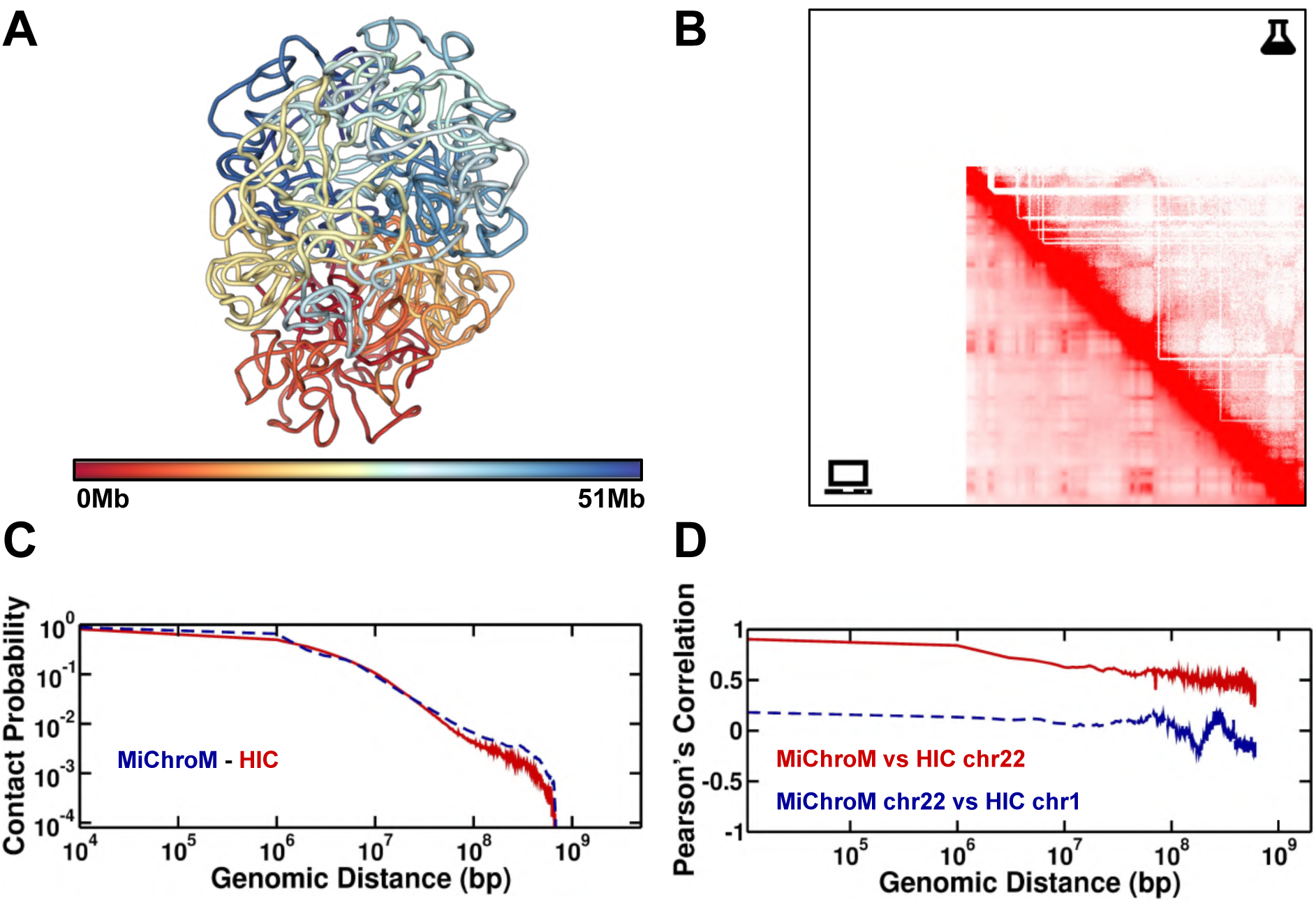
The structural ensemble of chromosome 22, cell line A549, generated by simulations using the MiChroM+MEGABASE pipeline. A - Three-dimensional representative structure colored by index, red to blue. B - Comparison between Hi-C maps obtained from wet-lab experiments (top) and *in silico* (bottom). The Hi-C maps predicted from simulations have been sampled from an ensemble of 500 thousand structures. C - Contact probability as a function of the genomic distance. The solid red line is the curve extracted from the experimental data. The dashed blue line is the data obtained from the *in silico* HiC. D - Pearson’s correlation between experimental and simulated Hi-C maps of chromosome 22 as a function of the genomic distance, shown by the solid red line. As a term of comparison the dashed blue line shows the correlation between Hi-C maps of different chromosomes.

